# Zero-shot evaluations expose structured generalization limits in predictive models of the brain

**DOI:** 10.64898/2026.09.15.751562

**Authors:** Ruolin Wang, Mayukh Deb, Alex Abate, Alish Dipani, Kushal Dudipala, Sanjana Chillarege, Kruthik Ravikanti, Yuxuan Li, Haider Al-Tahan, Ranjani Koushik, Elizabeth Mieczkowski, Herrick Fung, Nancy Kanwisher, N. Apurva Ratan Murty

**Affiliations:** Center of Excellence in Computational Cognition, Georgia Tech; School of Psychological and Brain Sciences, Georgia Tech; Harvard Medical School; Department of Computer Science, Princeton University; McGovern Institute for Brain Research, MIT; Department of Brain and Cognitive Sciences, Massachusetts Institute of Technology, MIT; The Center for Brains, Minds and Machines, MIT

**Author notes:** Correspondence: Ruolin Wang, N. Apurva Ratan Murty. These authors contributed equally (Ruolin Wang, Mayukh Deb).

## Abstract

Artificial neural networks (ANNs) can predict neural responses to natural images with remarkable accuracy, making them promising models of human vision. However, prediction scores alone reveal little about what computations these models have captured or how well they generalize beyond the images on which they are trained. Here, we introduce a principled zero-shot model evaluation framework in which ANN-based encoding models are frozen before being tested on not just held out images but entirely new datasets. Our strongest tests repurpose decades of cognitive neuroscience experiments as diagnostic benchmarks for identifying which computations current models capture and where they fail. To implement this framework we built encoding models of human category-selective regions (FFA, PPA, and EBA) using several pretrained ANNs and evaluated these fixed models across 7 independent fMRI datasets and 31 cognitive experiments from 10 published studies. This framework revealed three broad patterns. First, zero-shot evaluations exposed systematic limits to model predictivity that were largely hidden by standard within-dataset cross-validation. These limits were structured varying across brain regions and model classes and depending strongly on the similarity between the images used to build and test the models. Second, current models reproduced some, but not all, classical findings from cognitive neuroscience, providing a diagnosis of the computations that current models have yet to capture. Finally, models that performed well on prediction tests also tended to succeed at reproducing cognitive neuroscience findings, suggesting that prediction and explanation are closely linked. Unlike prediction scores however, cognitive tests help diagnose where and why models fail. Together, the zero-shot evaluation framework provides a scalable and unified approach for evaluating brain models across prediction and cognitive neuroscience tests, which can help us better understand what current models capture and what still remains to be explained.

## Introduction

Artificial neural network (ANN)-based models are routinely touted as leading models of primate vision [16, 21, 32, 40, 41, 65, 70, 73, 75, 76, 85, 86]. Indeed, these are the first image-computable models capable of predicting a substantial fraction of variance in neural responses. But the value of any scientific model depends on its ability to generalize beyond the observations used to construct it [61, 68]. Yet whether current ANN-based encoding models generalize in this broader sense remains largely untested. Current ANN-based encoding models are evaluated almost exclusively on randomly selected held-out images from the same fMRI datasets used to establish the brain mapping. While these within-dataset (cross-validated) evaluations quantify model performance on unseen images drawn from the same dataset, they provide only a limited assessment of how well models generalize. A more demanding test is whether the same models generalize to new subjects across independent datasets, and to the carefully controlled stimulus manipulations that have shaped decades of cognitive neuroscience and psychology. Cognitive experiments are particularly diagnostic because they use carefully designed, hypothesis-driven stimulus manipulations to dissociate visual properties that are otherwise correlated in natural image datasets (like Imagenet([18])). Here, we evaluate ANN models of the FFA, PPA, and EBA under a strict zero-shot framework that freezes every aspect of the encoding model before testing (**Fig.1a**). We first ask whether these models generalize across 7 independent fMRI datasets without retraining, and then whether they reproduce the neural responses observed across 31 cognitive neuroscience experiments. This work provides the first large-scale quantitative assessment of model generalization across independent datasets and cognitive neuroscience experiments, revealing structured limits that remain hidden in conventional within-dataset evaluations. By exposing both the successes and failures of current models, these evaluations provide a roadmap for developing faithful *in silico* models of the human visual system that can predict new experiments, reveal missing computations, and ultimately serve as reliable digital twins for vision research.

**Figure 1.**
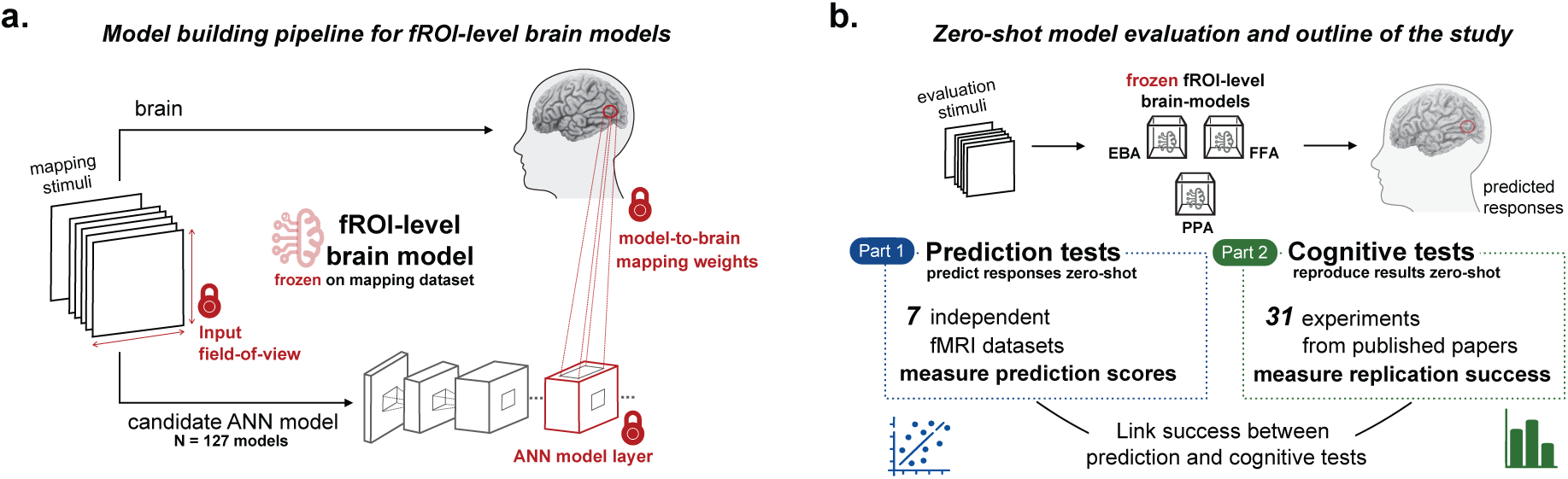
Zero-shot framework. **a. Construction of frozen fROI-level encoding models.** Functional regions of interest (FFA, PPA, and EBA) are localized in individual participants, and voxel-wise encoding models are built by mapping layer features from candidate ANNs to fMRI responses measured on a mapping dataset. Three components are fixed during model construction (red locks): the input field-of-view, the ANN layer with the best prediction, and the model-to-brain mapping weights. Once these choices are made, the encoding model is frozen and no further dataset-specific fitting is permitted. **b. Zero-shot evaluation framework and outline of the study.** Frozen fROI-level models are evaluated on independent fMRI datasets and experiments. Part 1 quantifies zero-shot *prediction* performance across public fMRI datasets, allowing us to compare models and identify the factors that determine prediction success. Part 2 repurposes classic cognitive neuroscience findings as zero-shot tests. We ask whether the same fixed models reproduce published findings to diagnose the computations they capture and those they fail to capture. Finally, we relate performance on prediction and cognitive tests to examine whether these complementary evaluations reflect a common notion of model quality.

Despite the growing adoption of ANNs in vision neuroscience, fundamental questions about their ability to generalize still limit their broader scientific acceptance [6, 29, 33, 44, 52, 72, 83, 88]. First, it is unclear if the predictions made by ANN-based encoding models generalize to new subjects and stimuli without requiring the models to be retrained each time new data are collected. This concern is often dismissed as a practical limitation. Neural data are often scarce, collecting new data is resource intensive, and it is almost impossible to identify the same neuron/voxel in a new individual. It is therefore a real possibility that current models overfit, and do not generalize beyond the dataset they were trained on [6, 22, 49, 64, 83]. A second challenge is that ANN-based brain models are often difficult for the broader neuroscience community to access, reproduce, and evaluate on independent datasets [30]. Consequently, the same brain models are rarely subjected to new datasets or unseen experimental paradigms. This makes it difficult to determine where current models succeed, where they fail, and what the structured limits of their generalization are.

But perhaps the strongest criticism is that the prediction success of ANN models has not made contact with the rich body of hypothesis-driven cognitive neuroscience experiments based on carefully controlled stimulus manipulations (‘*severe tests*’ [6]) developed to distinguish between competing theories of vision [35]. For example, hypotheses about the neural basis of face recognition have been tested on controlled stimulus manipulations like inverted faces [25, 87], face pareidolia [82], and faces in which the internal or external features of the faces have been scrambled [39] (among other manipulations). Each manipulated stimulus set evaluates a specific cognitive hypothesis about the neural mechanisms of face processing (e.g., the face inversion effect, holistic processing, etc.). These stimuli are not part of any standard ANN training dataset and often fall outside the natural image manifold altogether. ANN models of neural responses have rarely been evaluated against this body of hypothesis-driven evidence, leaving it unclear whether they capture the same computational principles of the visual cortex [7, 22, 29, 50].

To address these concerns, we propose a new model evaluation framework that tests computational models without retraining, across independent datasets and against hypothesis-driven experiments that have been central to cognitive neuroscience and psychology. The key conceptual advance is the scaled-up zero-shot (across-dataset) evaluation of brain-mapped models against previously published findings that *also* includes neural responses to highly manipulated non-naturalistic images. We argue here that the key test for a model is whether it generalizes beyond the data on which it was trained. Critically, all models are evaluated after removing all experimenter degrees of freedom (i.e. with exactly zero free parameters, or in a ‘*zero-shot*’ manner). We demonstrate this framework in three extensively studied (category-selective) regions of the high-level human visual cortex: the fusiform face area (FFA), the extrastriate body area (EBA) and the parahippocampal place area (PPA). These regions are ideal test cases because 1) they can be consistently identified in every participant using well-established robust localizers [20], and 2) have been extensively studied using stimulus manipulations to test their functional and computational roles. Applying this framework across two complementary tests, we show that the predictivity of ANN models has systematic limits. Prediction tests ask how faithfully a fixed model predicts neural responses to unseen data across datasets, whereas cognitive tests ask whether the same model reproduces published neural effects using the original experimental stimuli. Across both tests, model performance declined approximately linearly as evaluation stimuli become more distant from the mapping images in mid-level perceptual space. Generalization also varies systematically across brain regions. Models often predicted *what* category-selective regions respond to, but not *how* they represent visual information, with the largest gaps in body-selective regions. Together, this work brings predictive models into contact with hypothesis-driven cognitive neuroscience to reveal the limits of their generalization, and provides a framework for measuring when, where, and how neural predictions fail beyond the data used to build them.

## Results

### The zero-shot evaluation framework

Our modeling framework builds on the functional-ROI approach in cognitive neuroscience [24, 37, 57, 71]. We evaluate encoding models on functionally defined category-selective brain region across datasets, experiments, and subjects. The framework proceeds in four steps. First, we measure responses to a set of mapping stimuli in the FFA, PPA, and EBA, identified in each participant using standard fMRI localizers. Second, we build voxel-wise encoding models for each candidate ANN by learning a linear mapping from the optimal intermediate ANN-layer features to brain (voxel) responses [16, 63, 86]. Building an encoding model involves three key experimenter choices that are made once and then fixed: the pixels-to-visual-angle mapping (degrees of visual angle-to-pixel conversion for ANNs), the ANN layer corresponding to the best model fits, and the specific mapping weights from ANN features to fMRI responses (schematized as locks in **Fig.1a**). Once these choices are made, the model is frozen. Third, stimuli from entirely new fMRI datasets are passed through the frozen model to generate model-predicted responses without any further layer-selection or dataset-specific refitting. Finally, these zero-shot predictions are compared with the actual measured experimental data. All key evaluations in this paper follow this procedure and are therefore strictly *zero-shot*.

The Results are organized into two parts (**Fig.1b**). Part 1 focuses on *prediction tests*. We compare our zero-shot (across-dataset) framework to the standard (within-dataset) cross-validation approach (the dominant standard in NeuroAI [14, 16, 74, 75]) and ask how well current ANN-based models predict neural responses across multiple datasets under zero-shot setting. We also ask what determines success at prediction: which datasets and brain regions are hardest to predict, what makes a dataset easy or hard for models to predict, and which model architectures generalize best. Part 2 focuses on *cognitive (neuroscience) tests*. We test whether the same models reproduce classic findings from cognitive neuroscience experiments using carefully controlled stimulus manipulations. We ask which experimental effects models capture and which they consistently miss, what makes a cognitive neuroscience test experiment easy or hard to replicate, which brain regions and models do best, and whether success on prediction tests is systematically related to success on cognitive neuroscience tests.

### Part 1. Zero-shot prediction tests expose the limits of current ANN-based encoding models

#### 1.1 Predicting NSD responses zero-shot

We begin by describing how we evaluate models on the Natural Scenes Dataset (NSD; [1]) entirely zero-shot. NSD is a widely used, high-quality 7T fMRI dataset in which participants viewed thousands of naturalistic images. Our first step was to establish model prediction ceiling (the maximum achievable predictivity given data reliability) and the prediction floor (the performance of models with no high-level visual representations) as reference points for every dataset. We defined the ceiling as the consistency of responses across participants (NeuroAI Turing Test, [11, 12, 26, 80]. For univariate analyses, we computed pairwise Pearson correlations of ROI-averaged fMRI responses across the 1,000 images shared by four NSD participants (^4^ = 6 pairs). Inter-subject correlations were high (eg. EBA: mean *R* = 0.75, *P* < 0.00001; **Fig.2a**, *left*), consistent with the known reliability of the NSD dataset. For multivariate analyses, we computed Spearman rank correlations between RDMs across subject pairs. Pairwise RDM correlations were also substantial for the NSD (eg. EBA: mean *R* = 0.52*, P <* 0.00001; **Fig.2b**, *left*). The prediction floor was estimated using three baselines: a pixel model, models approximating V1 responses (VOneNet [17]), and several untrained ANNs (see Methods).

**Figure 2.**
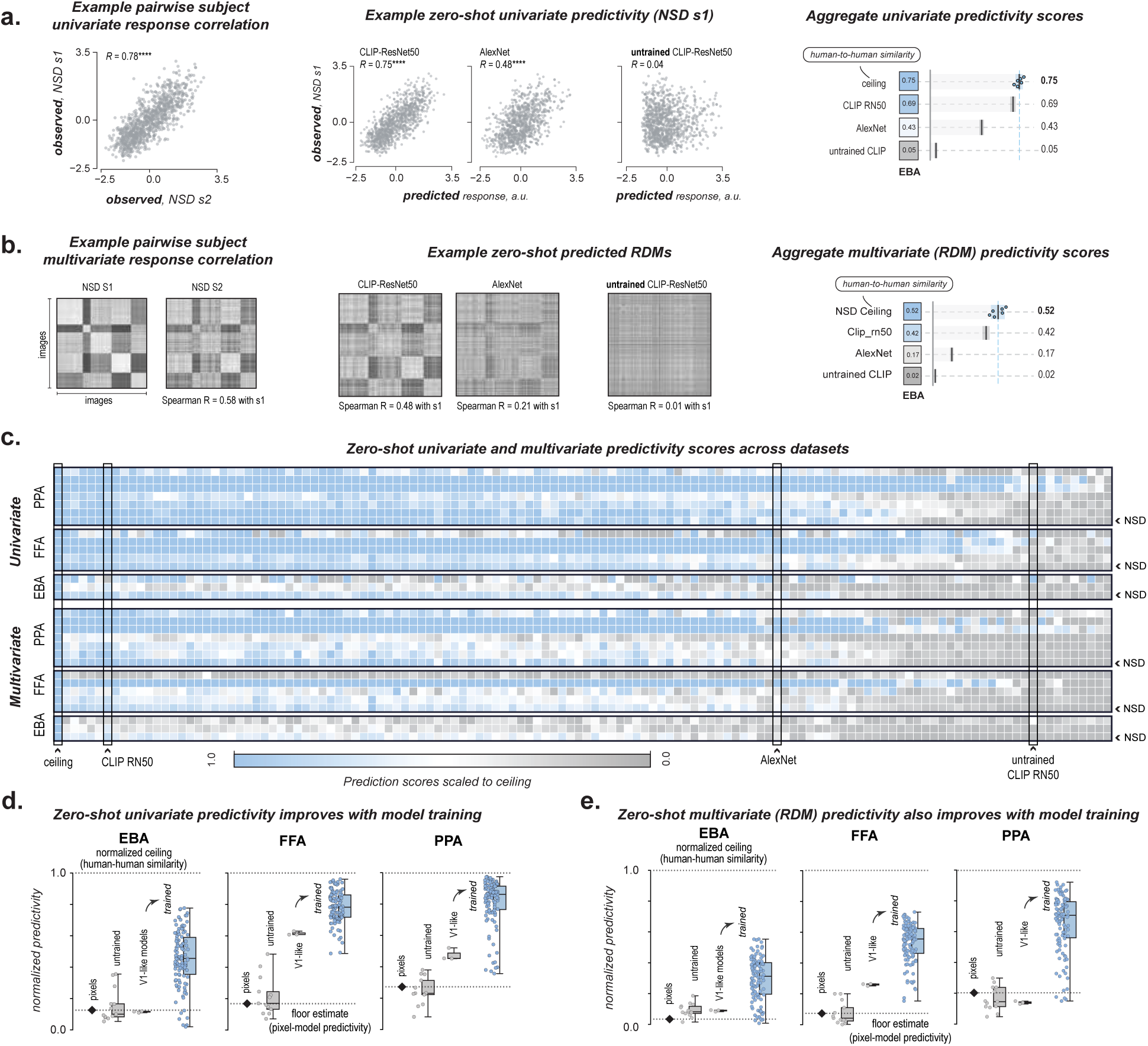
Zero-shot model evaluations on fMRI datasets. **a. Zero-shot univariate evaluation for NSD EBA**. *Left*, Scatterplot comparing voxel-averaged EBA responses from two NSD participants across shared images, illustrating high inter-subject similarity. *Middle*, Example zero-shot predictions from CLIP-ResNet50, AlexNet, and an untrained CLIP-ResNet50 model plotted against measured responses from NSD subject 1. *Right*, Aggregate subject-averaged univariate prediction scores for EBA across the three models. **b. Zero-shot multivariate evaluation for NSD EBA.** *Left*, Representational dissimilarity matrices (RDMs) computed from two NSD participants. *Middle*, Example zero-shot RDM predictions from CLIP-ResNet50, AlexNet, and an untrained CLIP-ResNet50 model compared with the measured RDM from NSD subject 1. *Right*, Subject-averaged multivariate (RDM) prediction scores for EBA across all evaluated models. **c. Zero-shot prediction across datasets.** Subject-averaged univariate (top) and multivariate (bottom) prediction scores for PPA, FFA, and EBA across all models and fMRI datasets. Models are ordered by decreasing prediction performance. Scores are normalized by the human-to-human similarity ceiling, allowing comparison across datasets. Each column corresponds to one ANN model and each row to one brain region and dataset. Only representative models are labeled; complete results for all 127 models are shown in Supplementary **Fig.S1**. **d. Zero-shot univariate prediction improves with model training.** Normalized univariate prediction scores for EBA, FFA, and PPA comparing pixel-level predictions, untrained models, VOne-based models, and trained ANN models. Each point represents one ANN model; boxes summarize the distribution across models. Dashed lines indicate the human-to-human similarity ceiling (top) and pixel-level prediction baseline (bottom). **e. Zero-shot multivariate (RDM) prediction also improves with model training.** Same analysis as in **d**, using multivariate (RDM) prediction scores.

We then asked how well ANN-based models predict NSD responses zero-shot. We trained voxel-wise encoding models for FFA, PPA, and EBA using an independent mapping dataset with only 185 images (henceforth Murty185 dataset [63], and resulting models Murty185-mapped models) previous shown to support reliable model-to-brain mappings in these regions. In this case, we fixed the model parameters based on Murty185, and evaluated each model on NSD without any NSD-specific model fitting (**Fig.1**), comparing both univariate (voxel-averaged) and multivariate (RDM-based) predictions. **Fig.2a** (*middle*) shows example scatterplots for three representative ANN models evaluated on EBA responses from the first NSD subject. ResNet50-CLIP achieved high predictive accuracy (*N* = 1000*, R* = 0.75*, P <* 0.00001), AlexNet showed lower accuracy (*R* = 0.48*, P <* 0.00001), and an untrained model performed near floor (*R* = 0.04*, P* = 0.17). The same pattern held for multivariate analyses (**Fig.2b**, *middle*, *right*). ResNet50-CLIP showed the strongest match with human multivariate representational structure (*R* = 0.48*, P <* 0.00001; inter-subject ceiling = 0.52), AlexNet was weaker (Spearman *R* = 0.21*, P <* 0.00001), and an untrained model showed near-zero prediction (*R* = 0.01*, P <* 0.00001). These examples show that it is possible to predict responses to NSD images entirely zero-shot without any retraining on NSD-specific images.

### 1.2 Scaling up zero-shot evaluation across models, datasets and brain regions

Next, we scaled our model evaluation process to 127 leading ANN models spanning convolutional networks, vision transformers, vision-language models, self-supervised architectures, and models incorporating brain-like constraints such as recurrence and topography (see full list in Supplementary Tables S4 and S5). We evaluated these models across six publicly available fMRI datasets that differ in stimuli, experimental design, scanner hardware, and participant populations. Datasets included images with natural images, (BOLD5000v2 [13], King [42], Bonner [5], Wardle [82]), short video stimuli (BMD [46]), and non-naturalistic synthetic images (NSD-synthetic [31]) across 3 category-selective brain regions (FFA, PPA, EBA). **Fig.2c** summarizes zero-shot univariate (*top*) and multivariate (*below*) prediction scores for every model, dataset, and all three brain regions (see Supplementary **Fig.S1** for a fully annotated version).

The results were consistent across brain regions (**Fig.2d,e**). Prediction accuracy increased from the pixel-model floor to V1-like models, and was highest for trained ANNs (on average). This pattern was consistent across all three brain regions (FFA, PPA, and EBA) and for both univariate (**Fig.2**d) and multivariate analyses (**Fig.2e**). To test whether the mapping dataset mattered for these analyses (**Section 1.3**), we trained a second set of encoding models on the shared 1000 images from NSD (henceforth NSD-mapped models). Trained NSD-mapped models also consistently outperformed the pixel-model floor and V1-like models (Supplementary **Fig.S3**).

### 1.3 Zero-shot evaluation enhances model differences and benefits from more mapping data

Having shown that ANN models can predict neural responses in a zero-shot setting, we next asked two questions. First, how does zero-shot evaluation compare with the standard within-dataset cross-validation used in NeuroAI? Second, does the choice of mapping dataset influence zero-shot prediction?

To compare zero-shot evaluation with the within-dataset cross-validation (the dominant standard in NeuroAI [16, 74]), we used the two sets of models built on independent mapping datasets and evaluated each set in two ways: on held-out images from the *same* dataset (standard within-dataset crossvalidation) and zero-shot from models based on the other dataset (across-dataset), which differed in stimuli, fMRI scanner, experimental design, and participants. The scatterplot in **Fig.3a** compares univariate and multivariate prediction scores under zero-shot (across-dataset, x-axis) and standard (within-dataset, y-axis) evaluations on the same images. We observed three patterns. First, models prediction scores were higher under standard evaluation than zero-shot evaluation (all points above x=y line, paired t-test P<0.0001) which suggests that standard cross-validation was still picking up some dataset-specific structure. Second, high performing models were more tightly clustered under standard evaluation (short edge of the rectangles) but became more separated under zero-shot evaluation (long edge of the rectangles). The interquartile range (IQR) more than doubled from 0.027 to 0.062 (2.3-fold increase) for univariate tests, and increased from 0.047 to 0.082 (1.8-fold increase) on multivariate tests. Third, lower-performing models shifted closer to the prediction floor under zero-shot evaluation. IQR scores decreased from 0.054 to 0.028 (0.5-fold) in the univariate analysis and from 0.034 to 0.014 (0.4-fold) in the multivariate analysis for the low-performing models. Despite these changes in absolute performance, the rank ordering of the 110 trained models was largely preserved (Spearman *ρ* = 0.81, P <0.00001). Reciprocal analyses using held-out Murty185 images yielded the same qualitative trends (data not shown). These results show that zero-shot evaluation provides a more discriminative assessment of model performance by increasing separation among high-performing models while pushing weaker models closer to the prediction floor.

**Figure 3.**
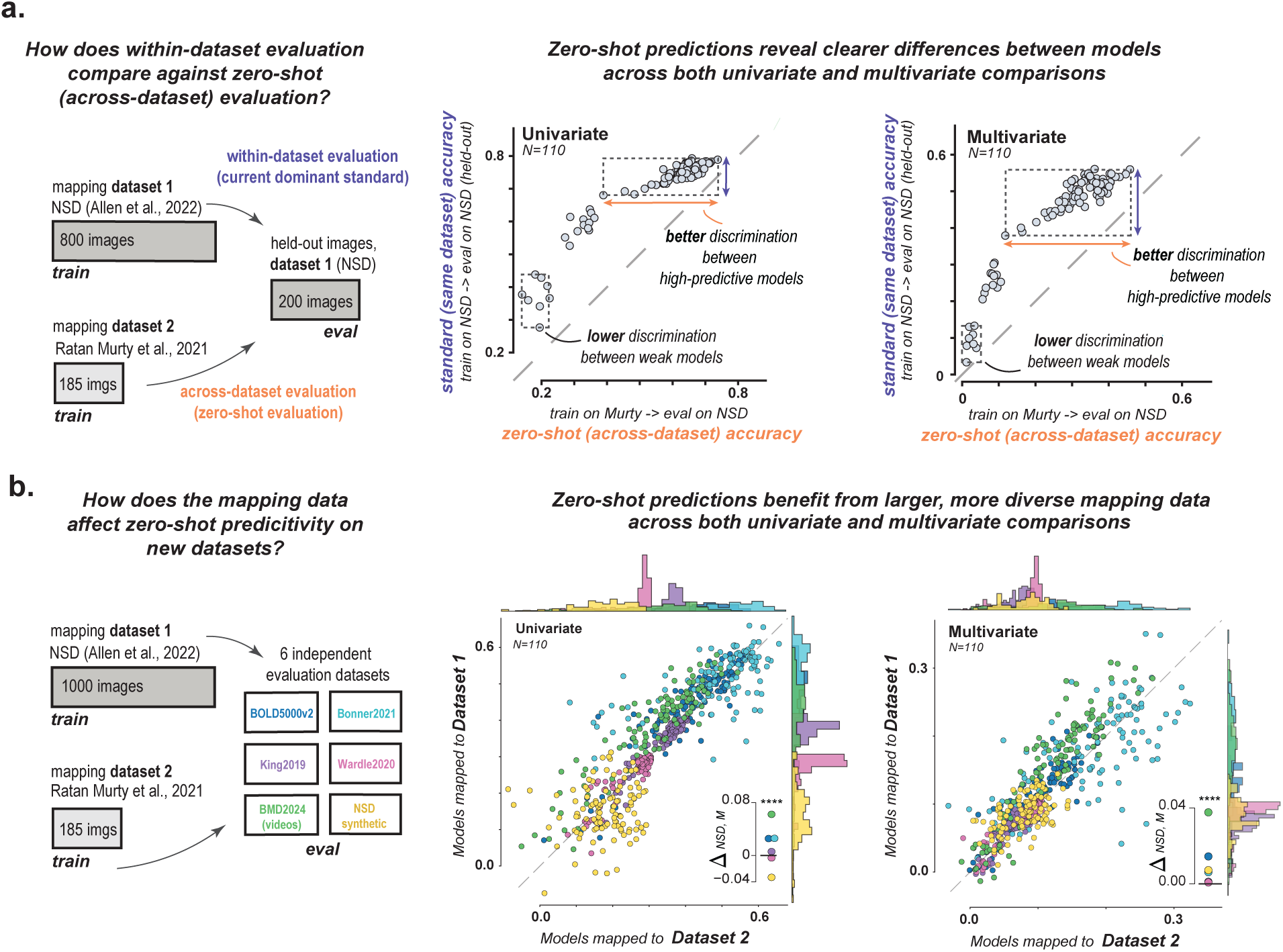
Zero-shot evaluation enhances model differences and benefits from larger, more diverse mapping datasets. **a. Zero-shot evaluation is more discriminative than within-dataset cross-validation.** *Left*, schematic comparing standard within-dataset evaluation with our across-dataset (zero-shot) framework. *Middle*, comparison of within-dataset and zero-shot univariate prediction scores across 110 trained ANN models. Zero-shot evaluation expands the dynamic range of prediction scores by separating high-performing models while driving poorly generalizing models toward chance performance. *Right*, same analysis using multivariate (RDM) prediction scores. **b. Zero-shot prediction benefits from larger, more diverse mapping datasets.** *Left*, schematic comparing encoding models built using the NSD (1000 shared images) and Murty185 (185 shared images) mapping datasets and evaluated on six independent fMRI datasets. *Middle*, comparison of zero-shot univariate prediction scores obtained using the two mapping datasets. Colors denote the evaluation dataset. *Inset*, median difference in zero-shot prediction scores (NSD−Murty185) across 110 trained models, illustrating the overall performance advantage of the larger NSD mapping dataset. *Right*, same analysis using multivariate (RDM) prediction scores.

We then asked how the choice of mapping dataset influences zero-shot prediction. To isolate the effect of the mapping data, we fixed the backbone ANN (architecture, training objective, and visual representations) and compared models mapped to the two mapping datasets (NSD-mapped models to the shared 1000 images and Murty-mapped models on the 185 shared images) on the same held-out datasets. Across 110 trained models, NSD-mapped models consistently achieved higher prediction scores across six independent evaluation datasets in both univariate and multivariate analyses (**Fig.3b**). Paired tests confirmed a modest but statistically significant advantage for the NSD-mapped models in both univariate (*t*(109) = 4.91, *p* = 3.20 10*^−^*^6^) and multivariate predictions (*t*(109) = 10.66*, p* = 1.31 10*^−^*^18^), with the largest gains for the Bold Moments (BMD) short-video dataset (univariate: *t*(109) = 13.76*, p* = 1.37 10*^−^*^25^; multivariate: *t*(109) = 11.70, *p* = 5.66 10*^−^*^21^; see **Tab.S1**). We therefore use the NSD-mapped models for the remaining analyses unless otherwise noted, and report the corresponding Murty185-mapped results in the Supplementary Materials. We return to the factors underlying this improvement below. Although the improvement was modest, it was consistent across models and evaluation datasets, indicating that the mapping dataset is an important component of zero-shot prediction performance.

### 1.4 Zero-shot evaluation reveals structured limits to neural prediction

Zero-shot prediction scores were not uniform across datasets or brain regions. Rather, prediction failures showed systematic structure, depending on both the stimulus domain being predicted and the target brain region. To characterize these limits, we quantified prediction difficulty as the normalized gap between average model performance and the estimated inter-subject ceiling (**Fig.4**). Results using a complementary Wasserstein distance metric between the full model and inter-subject score distributions are shown in Supplementary **Fig.S5** and lead to the same conclusions.

**Figure 4.**
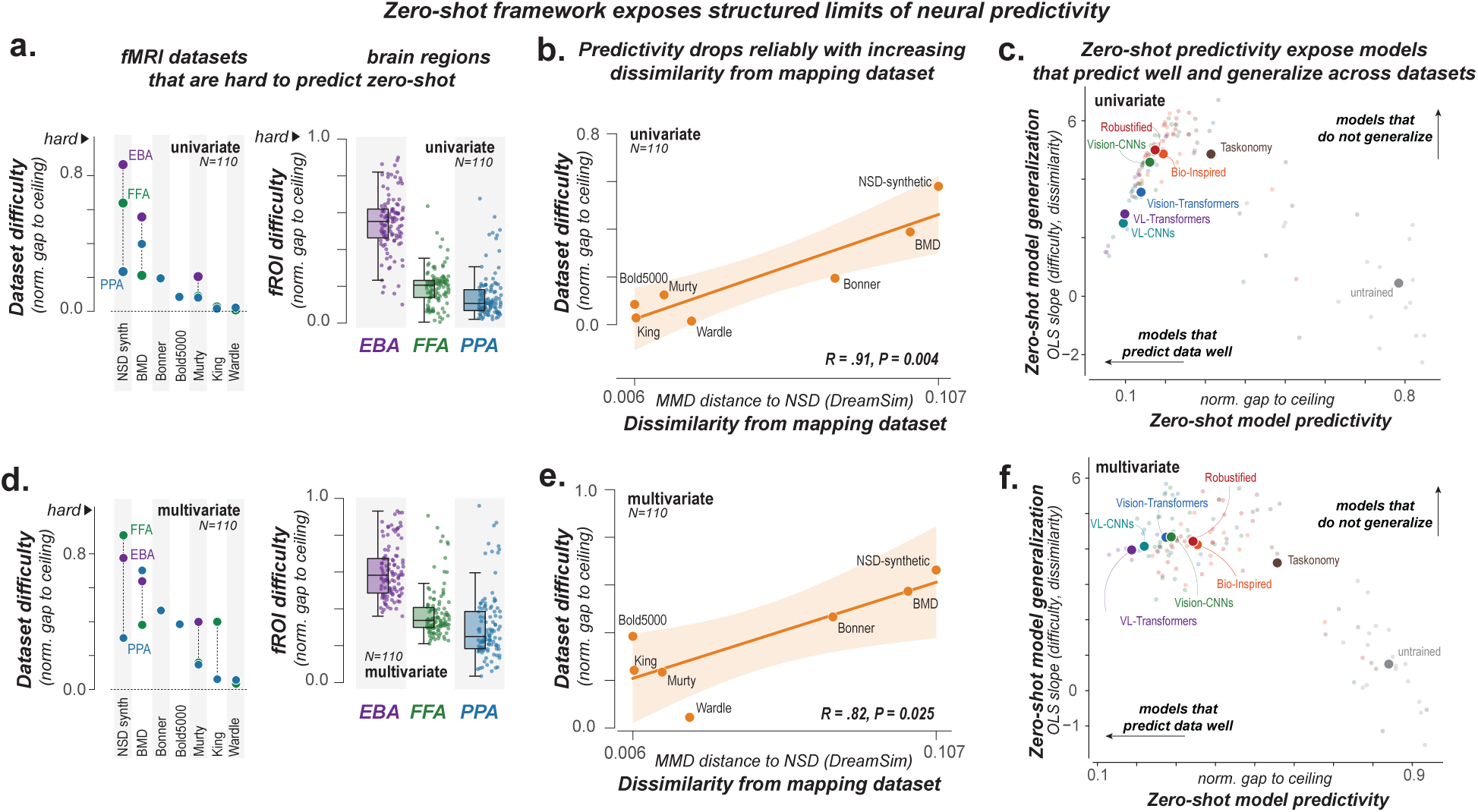
Zero-shot evaluation exposes structured limits to neural prediction. **a.** Prediction difficulty varies systematically across datasets and brain regions. *Left*, univariate prediction difficulty for each evaluated fMRI dataset, quantified as the normalized gap to the human-to-human similarity ceiling. Colors indicate the corresponding fROI. *Right*, fROI difficulty (y-axis) measured as the gap to ceiling across datasets for univariate data. **b.** Dataset difficulty increases with dissimilarity from the mapping dataset. Zero-shot univariate prediction difficulty plotted against the visual dissimilarity (DreamSim MMD) between each evaluation dataset and the mapping dataset (NSD). Prediction accuracy declined systematically as evaluation datasets became more dissimilar from the mapping images. **c.** Zero-shot evaluation identifies specific model families that both predict well and generalize well across datasets. Each point represents one ANN model. Zero-shot prediction performance (median normalized gap to ceiling, x-axis), versus zero-shot generalization across datasets (slope of prediction performance across evaluation datasets, y-axis). Models in the bottom-left simultaneously achieve strong prediction accuracy and strong generalization. **d.** Same analysis as in **a.** using multivariate (RDM) prediction scores. **e.** Same analysis as in **b.** using multivariate (RDM) prediction scores. **f.** Same analysis as in **c.** using multivariate (RDM) prediction scores.

Prediction difficulty varied substantially across datasets (**Fig.4a,d**). Out-of-distribution stimuli (NSD Synthetic; [31]) and natural videos (BOLD Moments; [46]) consistently showed the largest gaps to the inter-subject ceiling (mean across the three brain region univariate *R* = 0.58 and 0.39; multivariate *R* = 0.67 and 0.57), indicating that the out-of-domain images and video stimuli in these datasets are difficult to predict zero-shot. But why are some datasets easier to predict zero-shot than others? One idea is that prediction ability depends on how similar the evaluation dataset is to the mapping dataset. To test this idea, we quantified dataset similarity using maximum mean discrepancy (MMD), a kernel-based measure of the distance between image distributions. Because there is no single notion of image similarity, we computed MMD in three complementary feature spaces spanning different levels of visual processing: pixels (low-level image statistics), DreamSim [27] (mid-level perceptual similarity), AligNet [56] (high-level perceptual similarity). Datasets that were more similar to the mapping dataset were easier to predict. Among the three similarity measures, the mid-level perceptual distance DreamSim showed the strongest association with prediction difficulty (univariate *R* = .91*, p* = 0.004, multivariate *R* = .82*, p* = 0.025; **Fig.4b,e**), outperforming both pixel-based (univariate *R* = .29*, p* = 0.52, multivariate *R* = .51*, p* = 0.25) and higher-level (AligNet, univariate *R* = .32*, p* = 0.49, multivariate *R* = .26*, p* = 0.57) based measures, as shown in Supplementary **Fig.S6**. To summarize, zero-shot prediction declines systematically with perceptual distance from the mapping data, with mid-level perceptual similarity providing the strongest predictor of this decline.

Zero-shot predictions were also anatomically structured. Across nearly every evaluation, EBA exhibited the largest normalized gap to the ceiling (univariate median = 0.55, compared with 0.21 for FFA and 0.11 for PPA; multivariate median = 0.58, compared with 0.34 for FFA and 0.25 for PPA; **Fig.4a,d**), independent of training objective or mapping dataset. The same ordering was observed when restricting the analysis to the three datasets that included all three regions (NSD Synthetic, BOLD Moments, and Murty185). Thus zero-shot generalization is determined by both the stimulus domain and the target brain region. Prediction declines with mid-level perceptual distance from the mapping data, with the most pronounced gaps in EBA within our set.

### 1.5 Zero-shot generalization differs systematically across ANN models

Finally, we asked whether some classes of ANN models consistently generalize better than others under zero-shot evaluation, and whether models with stronger brain predictivity also generalize more robustly across increasingly dissimilar datasets. We grouped the 127 ANN models into eight families: vision-language transformers (VL-Transformer, *n* = 7; e.g., CLIP-ViT, BLIP-2, SigLIP), vision-language CNNs (VL-CNN, *n* = 3; e.g., CLIP-RN50), vision-only transformers (*n* = 11; e.g., DINOv2, BEiT), vision-only CNNs (*n* = 30; e.g., ResNets, VGGs), robustified CNNs (*n* = 27; adversarially and blur-trained variants), bio-inspired models (*n* = 15; CORnets, TDANNs, TopoNets), task-specific models (*n* = 17; Taskonomy), and untrained or pixel-level baselines (*n* = 17; Random, Pixels, VOne-Layer). For each model, we summarized overall predictive performance using the normalized prediction gap, defined as the difference between model performance and the human ceiling (lower values indicate better performance). To quantify robustness to dataset shifts, we computed the generalization slope, defined as the rate at which the prediction gap increased as evaluation datasets became increasingly dissimilar (based on DreamSim) from the mapping dataset (lower values indicate greater robustness). For visualization in **Fig.4c,f**, each model family is summarized by its median prediction gap.

Vision-language models achieved the smallest prediction gaps (VL-Transformer: 0.10 univariate, 0.19 multi-variate; VL-CNN: 0.10, 0.22), followed by vision-only transformers (0.14, 0.27) and vision-only CNNs (0.16, 0.29) (**Fig.4c,f**). Taskonomy models (0.31, 0.56) and untrained baselines (0.78, 0.84) performed substantially worse than all other trained model classes (Mann–Whitney *U*, both *p <* 10*^−^*^9^). Although vision-language models exhibited the strongest overall brain predictivity, this advantage should be interpreted cautiously because these models are typically trained on substantially larger image collections than vision-only architectures.

Consistent with this possibility, among models trained on similarly large datasets (millions of images, log_10_(ImagesNum) [1.15, 4.00]), the difference between vision-language and vision-only models was no longer significant (univariate: prediction gap = 0.09 vs. 0.12, *n* = 9 vs. 6, Mann–Whitney *U* test, two-sided *p* = 0.27; multivariate: 0.19 vs. 0.23, *n* = 9 vs. 6, *p* = 0.15). Thus, vision-language and vision transformer models showed the strongest overall zero-shot brain predictivity, although the apparent advantage of vision-language models could not be separated from differences in training data scale.

We next asked whether the models that best predicted brain activity also generalized more robustly as evaluation datasets became dissimilar from the mapping dataset. We exclude untrained baselines from this analysis because their predictions were at floor across evaluation datasets and thus produce artificially shallow generalization slopes (trained models, *n* = 110: univariate median slope = 4.61, IQR 3.65 − 5.47; multivariate median = 4.17, IQR 3.81 − 4.68; untrained baselines, *n* = 17: univariate median = 0.45; multivariate median = 0.75; Mann–Whitney *U* test, both *p <* 10*^−^*^9^). Among trained models, smaller prediction gaps were associated with shallower generalization slopes in the univariate analysis (Spearman *ρ* = 0.53, *p <* 10*^−^*^8^), indicating that models with stronger brain predictivity retained their performance better as evaluation datasets became increasingly dissimilar from the mapping dataset. Consistent with this relationship, models in the best-performing quartile showed shallower generalization slopes than the remaining models (median = 3.40 vs. 4.89; Mann–Whitney *U* test, *p <* 10*^−^*^9^). This relationship was absent in the multivariate analysis. Prediction gap and generalization slope were not significantly associated (*ρ* = −0.05, *p* = 0.61), and models in the best-performing quartile showed no clear generalization advantage over the remaining models (median slope = 4.03 vs. 4.29; *p* = 0.11). Thus, stronger univariate brain predictivity was associated with greater robustness to dataset shift, whereas stronger multivariate predictivity did not confer a comparable generalization advantage.

At the individual model level, BLIP-2, KOSMOS-2, Nomic, and EVA-2 ranked among the top five models for brain predictivity in both univariate and multivariate evaluations. In the univariate evaluation, these models also exhibited some of the shallowest generalization slopes among high-predictivity models, combining strong brain predictivity with robust cross-dataset generalization (BLIP-2: prediction gap = 0.05, slope = 1.51; KOSMOS-2: 0.05, 1.49; Nomic: 0.06, 1.72; EVA-2: 0.06, 1.37). In the multivariate evaluation, the same models remained among the most predictive, but their generalization slopes were comparable to those of other high-predictivity models (BLIP-2: prediction gap = 0.13, generalization slope = 3.84; KOSMOS-2: 0.15, 3.97; Nomic: 0.19, 3.95; EVA-2: 0.19, 4.08), consistent with the weaker relationship between predictivity and robustness observed across models.

### Part 2: Zero-shot cognitive tests reveal the functional properties captured by ANN-based encoding models

A model that predicts responses to natural images may still leave open what features of those images are driving the prediction. Cognitive neuroscience addresses this problem with controlled stimulus manipulations that separate features that are correlated in natural images. These experiments provide especially strong tests of brain models. If a model captures the relevant functional properties of a region, it should reproduce the response patterns elicited by brain to these manipulations. We therefore use these experiments to ask which established findings current models recover, which brain regions and model classes are best captured, and how success on these “cognitive (neuroscience) tests” relates to quantitative neural prediction (Part 1).

### 2.1 Running cognitive tests entirely zero-shot

We shortlisted 31 experiments from 10 published studies that met three criteria: (1) they reported responses from individually localized FFA, PPA, or EBA; (2) the original stimuli were publicly available; and (3) they tested a specific hypothesis about visual cortex function, typically using carefully controlled stimulus comparisons or manipulations. We then ran each experiment entirely zero-shot on our ANN-based encoding models by presenting the original stimuli and generating predicted neural responses without any dataset-specific fitting. Because these experiments were designed to test hypotheses about the brain rather than to evaluate artificial neural networks, they provide particularly diagnostic tests of whether current models reproduce established findings from cognitive neuroscience. **Fig.5a,b** illustrate two representative experiments. **Fig.5a** shows an experiment testing holistic face processing in the FFA and EBA [39] in which responses were measured to faces with intact or scrambled internal and external features. **Fig.5b** shows an experiment relating representational geometry in FFA to human visual search behavior [15], in which pairwise neural dissimilarities were compared with visual search reaction times. These examples show the range of hypothesis-driven experimental designs included in our evaluation.

**Figure 5.**
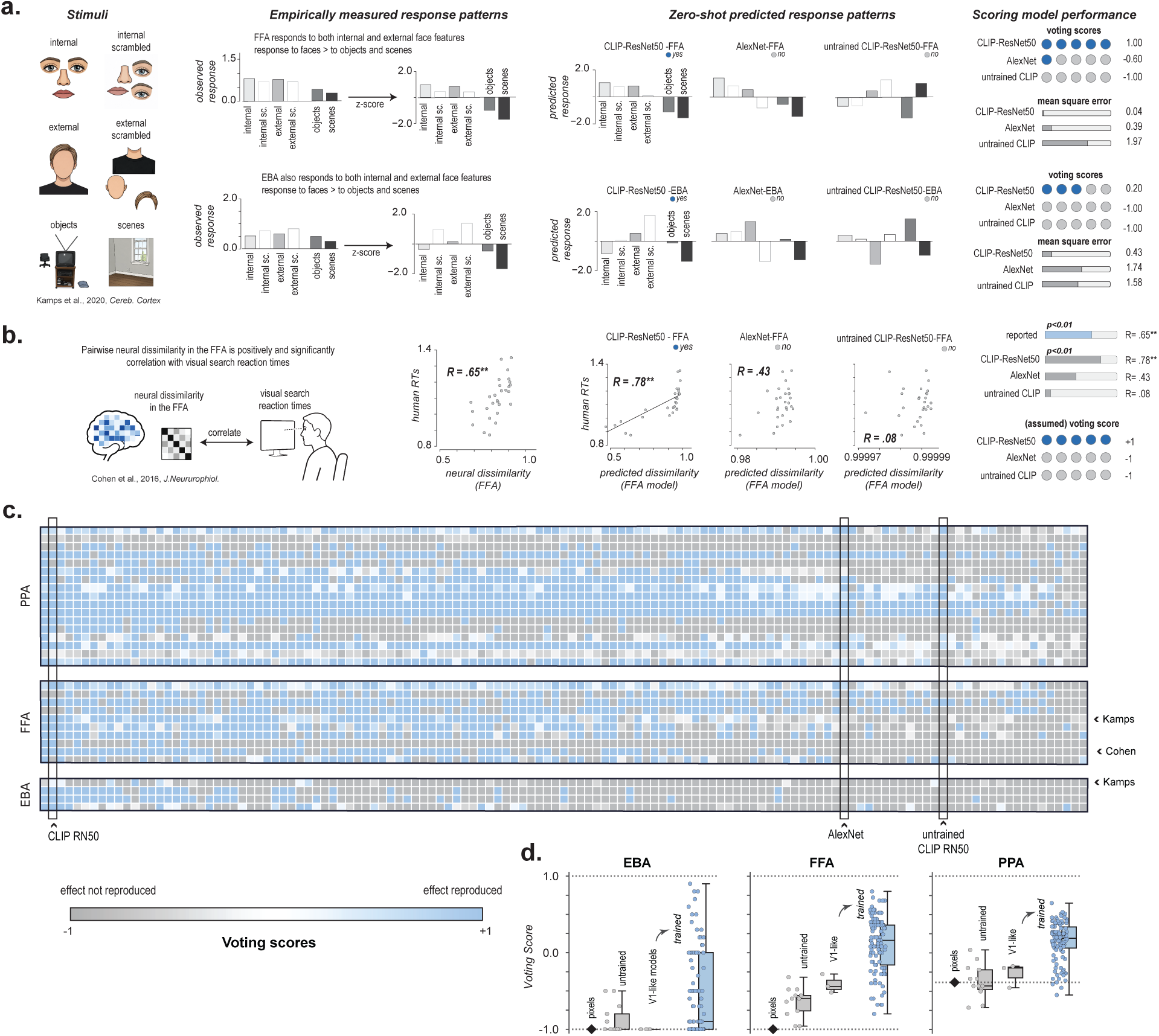
Zero-shot model evaluations on cognitive tests. **a. Zero-shot evaluations on Kamps et al 2020**. From left to right, *Stimuli*: the original experiment setup; *Empirically measured response patterns*: the result pattern reported in paper figure, after z-score, and comparison with the pattern replicated by three representative models: CLIP-ResNet50, AlexNet, and untrained CLIP-ResNet50; *Scoring model performance*: Human raters judged whether each model reproduced the experimental result; the voting score is defined as *p*_yes_ *− p*_no_ *∈* [*−*1, 1](all yes *→* +1, all no *→ −*1). As a complementary metric, mean squared error (MSE) compares each model’s z-scored predicted condition profile to the paper’s z-scored fROI response profile. **b. Zero-shot evaluations on Cohen et al 2016.** Cohen asks whether cross-category neural similarity correlates with the category-pair reaction-time. Based on the paper-specific statistical criteria, here the voting score is not from human raters, but assumed to be +1 if this correlation is positive and significant (*p <* 0.01), and *−*1 otherwise. **c. Zero-shot cognitive tests across experiments.** Voting scores for PPA, FFA, and EBA across all models and 31 experiments from 10 papers. Models are ordered by decreasing prediction performance. Each column corresponds to one ANN model and each row to one brain region and experiment. Only representative models are labeled; complete results for all 127 models are shown in Supplementary **Fig.S2**. **d. Zero-shot cognitive tests performance (voting score) improves with model training.** Voting scores for EBA, FFA, and PPA comparing pixel-level predictions, untrained models, VOne-based models, and trained ANN models. Each point represents one ANN model; boxes summarize the distribution across models. Murty-mapped models result is shown in Supplementary **Fig.S4**.

For each of the 31 experiments, we asked whether a model reproduced the central finding reported in the original study. We summarized this outcome with a common voting score (see Methods), which indicated whether the model captured the key experimental effect. The score was obtained in one of two ways, depending on the form of the original result. For experiments in which the conclusion depended on the pattern of responses (as in **Fig.5a**), each model under each mapping source (NSD or Murty) was independently judged by five of ten neuroscience researchers on whether the model reproduced the key experimental effect. **Fig.5a** shows the observed responses from the original experiment alongside predictions from three representative ANN models and the corresponding voting outcomes. Although this evaluation necessarily involves human judgment, agreement across raters was high (Spearman-Brown corrected split-half reliability: mean ± SD = 0.86±0.02). For experiments in which the original conclusion was defined by an explicit statistical test (as in **Fig.5b**) we applied the same statistical analysis to the model predictions and assigned the voting score (−1 or +1) directly from that result. **Fig.5b** similarly shows the relationship between model-predicted neural dissimilarities and observed visual search reaction times for three representative models.

We performed another independent validation of these voting-based evaluations. We digitized data from the published figures using WebPlotDigitizer [67] and computed the mean squared error (MSE) between the reported neural responses and model predictions. Voting scores were strongly associated with these independently derived MSE measures (*r*(125) = −0.91, *P <* 0.00001 for NSD-mapped models; *r*(125) = −0.89, *P <* 0.00001 for Murty-mapped models), indicating that models judged to reproduce the experimental findings also more closely matched the reported response patterns quantitatively.

Model rankings were also highly consistent across the two independently constructed mapping datasets (Spearman-rank correlation *ρ* = 0.76, *P <* 0.00001 between NSD-mapped and Murty-mapped models voting ranks, and *ρ* = 0.79, *P <* 0.00001 MSE ranks), although NSD-mapped models were modestly better among the top-performing models. Unless otherwise noted, subsequent analyses therefore use voting scores from the NSD-mapped models (analyses using Murty-mapped models are reported in the Supplementary **Fig.S7** and yield the same overall conclusions).

**Fig.5c** summarizes performance across all 31 experiments, 127 models, and three brain regions, providing an overview of which established cognitive effects are consistently captured by current models and which remain challenging. Trained ANNs reproduced more cognitive neuroscience findings than pixel and V1-like baselines (**Fig.5d**). We next examine which experimental effects current models reliably capture, which they systematically miss, and how these successes and failures vary across brain regions and model classes.

### 2.2 Zero-shot cognitive tests reveal structured limits to model generalization

The 31 cognitive experiments varied widely in how often their central findings were reproduced by current ANN models. Some effects were recovered by nearly every trained model, whereas others were missed almost uniformly. To understand this variation, we grouped the experiments according to the functional property they were designed to test and ordered them by replication success across models (**Fig.6a**). Broad category preferences and relatively coarse scene properties were among the easiest effects to reproduce. These included category selectivity [36], scene clutter [58], reachspace selectivity [36], face pareidolia [82], and scene content [36], for which many trained models recovered the expected response patterns.

**Figure 6.**
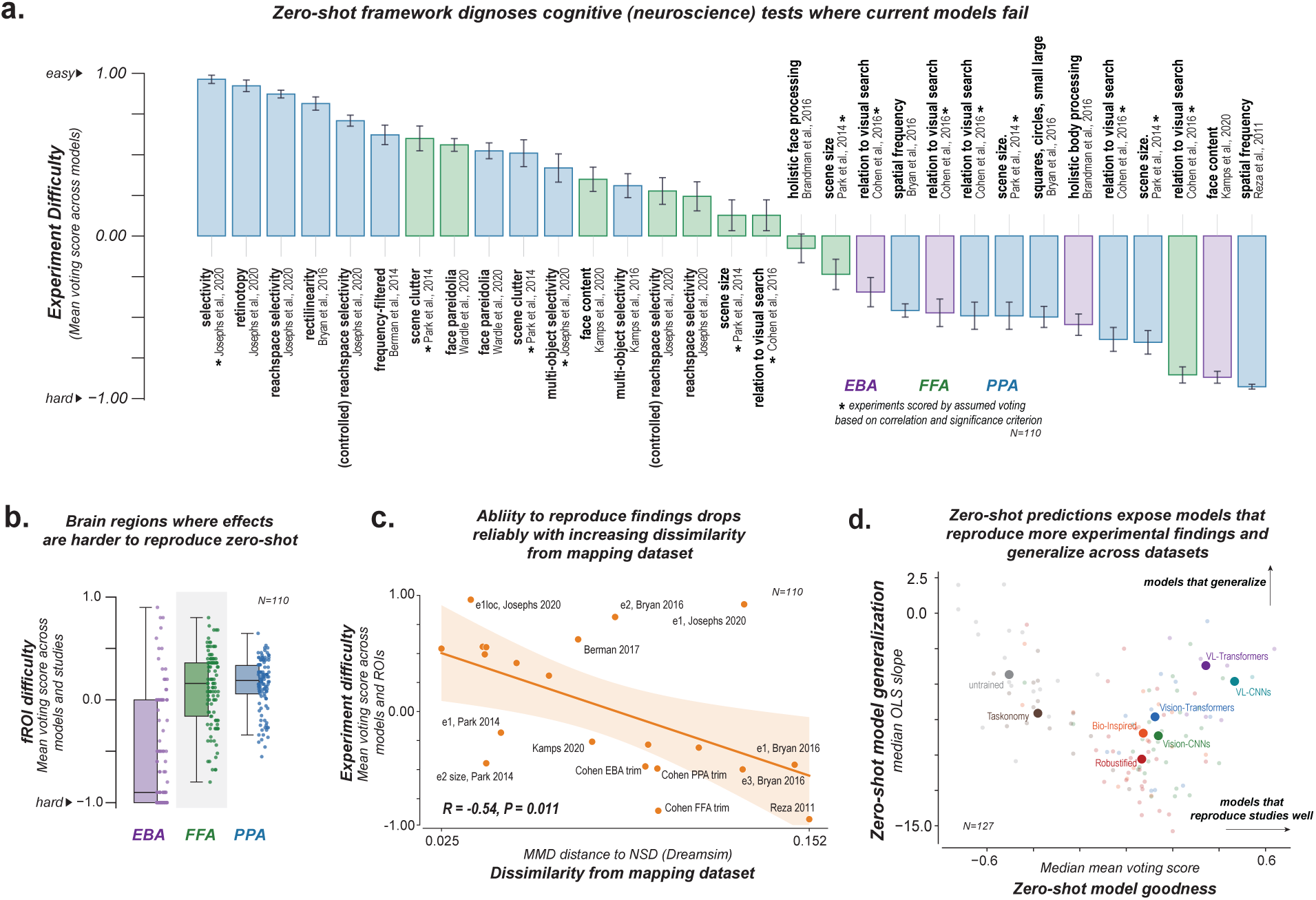
Understanding the challenges of model explanation on studies. **a.** Zero-shot evaluation identifies cognitive neuroscience findings that current models reproduce and those they consistently fail to capture. Thirty-one cognitive neuroscience experiments are ranked by replication success (y-axis; voting score), from easiest (left) to hardest (right). Colors indicate the corresponding category-selective region (EBA, FFA, or PPA). Positive scores indicate successful replication of the published finding, whereas negative scores indicate systematic failures. **b.** Brain regions differ in the difficulty of reproducing cognitive neuroscience findings. Replication success (voting score, y-axis) summarized across experiments for EBA, FFA, and PPA. Each point represents one model; boxes summarize the distribution across experiments. **c.** Experiments become harder to reproduce as they become more dissimilar from the mapping dataset. Replication success is plotted against the visual dissimilarity (DreamSim MMD) between each cognitive neuroscience experiment and the mapping dataset **d.** Models that generalize across datasets also reproduce more cognitive neuroscience findings. Each point represents one ANN model. The x-axis shows overall replication success across the 31 cognitive neuroscience experiments, and the y-axis quantifies zero-shot generalization across experiments (median OLS slope). Models in the upper-right reproduce the largest fraction of cognitive neuroscience findings and generalizing consistently across experiments.

Model failures were equally informative because they clustered around particular kinds of computations. Models rarely reproduced the relationship between neural representational geometry and human visual search behavior [15]. In these experiments, image pairs represented more similarly in FFA, PPA, or EBA were also harder for people to distinguish during visual search, resulting in longer reaction times. This link between neural similarity and behavior was rarely recovered from the model predictions in any of the three regions. Models also struggled with experiments probing holistic or configural processing of faces and bodies [8, 9, 39]. These studies asked whether responses depend on the spatial arrangement of features within a coherent face or body, for example by scrambling internal and external features or disrupting their normal configuration. Most models failed to reproduce these effects. Several classic response properties of scene-selective cortex were also difficult to recover, including sensitivity to spatial frequency and rectilinearity [3, 10, 62]. The experiments that current models failed to reproduce are therefore not a random subset of the literature. They disproportionately probe fine-grained representational geometry, configural processing, and specific visual features that go beyond broad category preferences.

The same pattern of structured failure appeared across brain regions. Across the 110 models, cognitive-test performance was substantially lower for EBA than for FFA or PPA (median voting score: EBA = −0.90, FFA = 0.16, PPA = 0.19, **Fig.6b**). This regional difference was highly reliable across models (Friedman *χ*^2^(2) = 104.85, *p <* 10*^−^*^22^). Pairwise comparisons confirmed lower performance for EBA than for either FFA or PPA (Wilcoxon signed-rank tests, both Holm-corrected *p <* 10*^−^*^15^), with no difference between FFA and PPA (*p* = .24). This difference was also evident within experiments that measured both EBA and FFA. Models performed worse in EBA for both Brandman and Yovel [9] (mean score 0.55 vs. 0.08, *p* = 4.0 10*^−^*^5^) and Kamps et al. [39] (−0.87 vs. +0.35, *p* = 4.4 10*^−^*^17^). The six statistically scored experiments from Cohen et al. [15] showed more variable regional differences and no consistent ordering. Thus, although fewer cognitive experiments were available for EBA overall, its poorer performance cannot be explained solely by differences in the experiments used to evaluate each region. Although fewer EBA experiments were available overall, this pattern mirrors the quantitative zero-shot results from Part 1 and identifies EBA as a consistent weakness of current ANN-based encoding models.

We then asked whether the difficulty of a cognitive experiment could be predicted from how far its stimuli departed from the images used to establish the encoding model. This analysis directly parallels the dataset-level result from Part 1. For each experiment, we measured the distance between its stimuli and the NSD mapping images in pixel space, DreamSim, and AlignNet, spanning low-level image statistics through increasingly abstract perceptual representations. Experiments whose stimuli were farther from the mapping images in DreamSim space were less likely to be reproduced by the models (*R* = −0.51, *P* = 0.027; **Fig.6c**). The corresponding relationships were weaker in pixel space (*R* = −0.15, *P* = 0.531) and AlignNet space (*R* = −0.31, *P* = 0.20). The cognitive tests therefore expose the same limits to generalization that emerged from the prediction tests. Models are most likely to fail when an experiment probes particular functional properties, when the target is EBA, and when the experimental stimuli lie farther from the images used to establish the brain mapping. These converging results suggest that model failures reflect systematic boundaries on what current ANN-based encoding models generalize to.

### 2.3 Zero-shot generalization on cognitive tests differs systematically across models

Across model families, replication success followed a clear hierarchy (Fig.6d). Vision-Language (VL) models achieved the highest voting scores overall (VL-CNN models median ≈ 0.47, VL-Transformers median ≈ 0.34). Vision-CNNs and Vision-Transformers formed a middle tier with more modest positive medians (median ≈ 0.13). Bio-Inspired and Robustified models sat lower still but remained slightly above zero (median ≈ 0.07). Taskonomy and floor models (untrained, pixel and V1-like) fell clearly below zero (medians 0.38 and 0.50), failing to replicate effects more than they succeeded. This ranking is consistent with the quantitative results from Part 1, reinforcing the conclusion that vision-language training confers advantages that show up whether models are evaluated on held-out neural data or on the targeted experimental effects that have shaped theories of high-level visual cortex.

Finally, we asked whether cognitive-test performance differed systematically across the model families (see Section 1.5), and whether models that reproduced more cognitive findings were also more robust across experiments with increasingly dissimilar stimuli. We quantified this robustness using the slope relating replication score to DreamSim distance from the NSD mapping images, with slopes closer to zero indicating greater robustness (as before). Vision-language models achieved the highest median replication scores (VL-CNN: 0.47; VL-Transformer: 0.34), followed by vision-only CNNs (0.14) and vision-only transformers (0.12) (**Fig.6d**). Taskonomy models (−0.38) and untrained baselines (−0.50) performed substantially worse than other trained model classes (Taskonomy vs other trained: Mann–Whitney *U* = 78, *p <* 10*^−^*^9^; untrained vs trained: *U* = 96, *p <* 10*^−^*^9^). We next asked whether models that reproduced more cognitive neuroscience findings also generalized more robustly as stimuli became more dissimilar from the mapping dataset. Among 110 trained models, replication performance was unrelated to generalization slope (Spearman *ρ* = −0.00, *p* = 0.99). Models in the highest-performing quartile showed only a modest shift toward shallower slopes than the remaining models (median = 7.07 vs. 8.68; Mann–Whitney *U* test, *p* = 0.05). Thus, stronger cognitive-test performance was not consistently associated with greater robustness to stimulus shift at the model level, echoing the weak relationship between predictivity and robustness in the multivariate analyses from Part 1. At the individual model level, Siglip2, CLIP-ResNet50, CLIP-ResNet101, Nomic, and Efficient-Net ranked among the top models for replication performance. Among high-scoring models, several also exhibited comparatively shallow generalization slopes, combining strong replication with more robust cross-experiment generalization (DinoV2-large: replication score = 0.35, slope = −0.57; Siglip2: replication score = 0.64, slope = −2.52; Blip2: replication score = 0.32, slope = −3.22; Nomic: replication score = 0.43, slope = −3.70). These results show that success on cognitive tests has two separable components: (1) how many established findings a model reproduces, and (2) how well the model prediction holds as stimuli get dissimilar to images in the mapping dataset (by a mid-level perceptual metric). The best predictive models perform well on both, but strong replication alone does not guarantee robust generalization.

### 2.4 Zero-shot predictivity and cognitive replication are tightly linked

Prediction tests and cognitive tests evaluate different aspects of a computational model. Prediction tests ask how accurately a model accounts for neural responses across a broad set of stimuli. Cognitive tests instead ask whether the model reproduces diagnostic response patterns observed in carefully controlled experiments designed to probe what a brain region represents or computes. Here we asked whether performance on these two forms of evaluation is related across models. Across models, quantitative prediction performance was strongly associated with performance on cognitive tests (**Fig.7a**). Models with higher cumulative prediction scores were more likely to reproduce the response patterns reported across the cognitive neuroscience experiments, for both univariate (Spearman *ρ* = 0.70, *p <* 0.0001) and multivariate prediction scores (*ρ* = 0.82, *p <* 0.0001). This relationship was not restricted to individual models. Model-family rankings were also largely preserved between prediction and cognitive tests (**Fig.7b**): model families that ranked highly on quantitative prediction tended to rank highly on the cognitive tests, whereas poorly predictive families, including Taskonomy and untrained models, remained among the lowest-ranked on both evaluations. Thus models that generalized better across independent datasets were also more likely to reproduce diagnostic effects from the cognitive neuroscience literature. This relationship has an important practical implication. Quantitative prediction tests (based on correlations) can be evaluated cheaply and at scale, providing a useful first-pass assessment of model quality, while cognitive tests provide a more targeted means of diagnosing which functional properties successful models have and have not captured.

**Figure 7.**
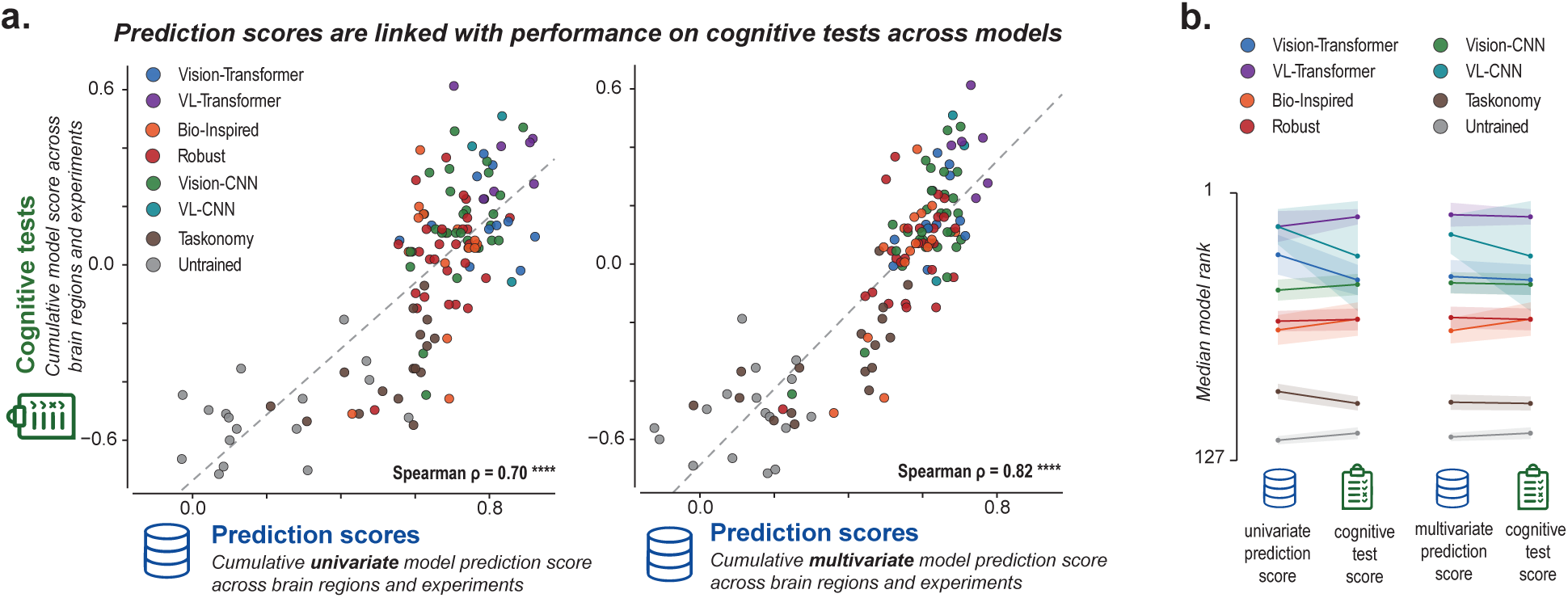
Prediction scores align with performance on cognitive tests. **a.** Scatterplots comparing each model’s cumulative cognitive-test score with its cumulative univariate prediction score (left) or multivariate prediction score (right), aggregated across brain regions and experiments. Each dot represents one model, and colors indicate model families grouped into broader model categories and the line indicates the least square line **b.** Model-family rankings are consistent across prediction scores and cognitive-test scores. For each model family, points show the median rank across models based on univariate prediction and cognitive-test performance (left), or multivariate prediction and cognitive-test performance (right). Lines connect the median ranks of the same model family across the two evaluation approaches, and shaded regions indicate the SEM across models within each family.

## Discussion

Scientific models are valuable only to the extent they generalize beyond the observations used to build them. Yet ANN-based predictive models are still evaluated primarily on randomly held-out stimuli from the same datasets used for model fitting, which makes it difficult to assess how broadly these models generalize or what computations they have actually learned [68, 69]. To address this problem, we introduced a scaled-up zero-shot evaluation framework that tests fixed fROI-level encoding models on independent datasets (**Sections 1.1 -1.3**) and cognitive neuroscience studies (**Section 2.1**) and applied it to the three canonical category-selective regions of human visual cortex (FFA, PPA, and EBA). Our results reveal the strengths and generalization limits of leading ANN-based encoding models. Predictions from many models generalized beyond the datasets used to build them, retaining substantial accuracy on entirely independent datasets without any additional fitting. Zero-shot evaluations also revealed systematic limits to this generalization (**Section 1.4**). Performance depended on the diversity of the mapping data, the brain region being predicted, and the perceptual similarity between mapping and evaluation stimuli. Cognitive experiments revealed a similar structure. Some classical effects were readily reproduced, whereas others were consistently missed, particularly when experiments used controlled stimulus manipulations to isolate visual properties that covary in natural images (**Sections 2.2 -2.3**). Together, these results show how broadly current ANN-based encoding models generalize, identify the functional properties and stimulus regimes that remain challenging, and place bounds on their current use as in silico “*digital twin*” models of visual cortex.

The zero-shot neural prediction tests in this work, extends the dominant benchmarking paradigm in NeuroAI [14, 74, 75]. At first glace, many ANN models do appear to retain substantial predictivity when tested on new fMRI data and participants without any refitting. Zero-shot evaluation nevertheless separated models more strongly than conventional within-dataset cross-validation (**Fig.3a**) [16, 63, 74, 77]. This pattern suggests that the apparent saturation of existing prediction benchmarks reflects, at least in part, the within-dataset evaluation strategy itself. Moving beyond the datasets used to map the brain models may expose more meaningful differences between candidate models. But prediction alone provides limited insight into what these models have captured and the gaps that remain.

The cognitive tests provide a more diagnostic assessment of these models by asking which established functional properties of category-selective cortex each model reproduces. We find that ANN models often accurately reproduce *what* these regions respond to, but do not match human response patterns with respect to *how* those brain regions process visual information (also consistent with [35, 76]. Matching broad response statistics was often sufficient to reproduce many well-known effects from the cognitive neuroscience literature, including category selectivity [36], sensitivity to scene clutter in the PPA [58], and face pareidolia [82] in the FFA. In contrast, models rarely produced the relationship between neural representational geometry and human visual search behavior[15]. Many models also consistently failed to reproduce effects that depend on integrating visual information into structured representations such as sensitivity to the relative arrangement of parts within faces and bodies in the EBA [9, 39] and spatial frequency tuning in the PPA [3, 62] (see also: [78]. This pattern reinforces the value of controlled manipulated stimulus tests for distinguishing models that perform similarly on natural images [4, 6, 23, 79].

A particularly important finding across both prediction and cognitive tests was a strong, approximately linear relationship between perceptual distance and model success (**Fig.4b,e** and **Fig.6c**). Models became progressively less accurate, and less likely to reproduce an experimental effect, as evaluation stimuli moved farther from the mapping images in mid-level perceptual space. This dependence was not weak for pixel distance and for the higher-level representations (AlignNet [56]) we tested, suggesting that the relevant boundary for the FFA, PPA, and EBA lies at an intermediate perceptual scale [27]. Whether the same scale governs generalization elsewhere in the brain remains an open question, particularly for regions encoding lower-level visual features (like V1) or more abstract conceptual information. At a fundamental level, the systematic fall-off with distance indicates that current ANN-based encoding models operate within an interpolation-like regime. The mapping images may define a domain within which model-to-brain correspondences are well constrained, with predictions becoming less accurate as stimuli beyond that domain. This idea has a practical consequence for model evaluation. Model performance should be tested with an explicit consideration of how far the test images are from the images used to build the mapping. Recent efforts to construct perceptually separated train test sets (as in MOSAIC [47] and LAION-fMRI) provide one way to make this distinction explicit.

Generalization limits were also structured across the brain regions we tested. We found that models fall consistently short at capturing body-selective responses of the EBA across both prediction tests (**Fig.4a,d** and **Fig.6b**) and cognitive tests. This pattern may reflect limitations in representing articulated and relational body structure [51], or in capturing the dynamics of bodies and social interactions [28, 48, 53]. Both possibilities remain open and point to important directions for future models.

Our results also speak directly to the recent debates over the relationship between prediction and explanation in NeuroAI [6, 29, 88]. Our results provide strong quantitative evidence to show that the distinction between the two may be less fundamental than recent debates suggest. Neural prediction and cognitive tests place different demands on a model. Prediction measures how accurately a model reproduces neural responses across many images, while cognitive tests ask whether it reproduces specific functional properties revealed by hypothesis-driven experiments [4, 33, 44]. In our analyses, we find that these two (seemingly opposing) notions of model quality converge to a striking degree (**Fig.7**). The models that produced the most accurate zero-shot predictions across independent datasets were also those that most faithfully reproduced cognitive experiments, despite some concerns that neural predictivity alone can under-constrain conclusions about brain-model correspondence [50, 77]. Thus, predictive accuracy becomes more informative when it is tested under sufficiently stringent conditions [12, 84] that require models to generalize beyond the data used to fit them. Cognitive tests then provide the complementary *diagnostic* step by telling us which functional properties successful models capture and where they still fail. Zero-shot evaluations provide a direct path from identifying models that predict the brain well (prediction tests) to determining the functional properties those models have actually learned about neural representation (cognitive tests).

On a related note, our findings were enabled by the zero-shot (across-dataset) evaluation framework, which we view as the primary methodological contribution of this work. Despite the rapid growth of open neuroimaging datasets [59, 60], most studies still evaluate models on held-out images within individual datasets. Our framework instead evaluates the same fixed set of models across independent datasets and decades of cognitive neuroscience studies, treating each as a new test of model generalization. This approach makes economical use of expensive fMRI data and experimental resources, by enabling researchers to subject models to many stringent tests without collecting a new dataset for every question. It also preserves the value of independent datasets as external tests of models, rather than repeatedly adapting models to each dataset [69]. Combining datasets and experiments can therefore expose model limitations that remain hidden within any single benchmark. Finally, the framework can be naturally extended to a new dynamic benchmark that can grow with the field as new datasets, experiments, and AI models become available.

Several limitations remain. First, we focused on category-selective regions because they provide a principled way to functionally align subjects and evaluate models entirely zero-shot [37, 71]. Extending the framework to additional functionally defined regions and ultimately to anatomically defined areas where correspondence across subjects is less direct is an important next step. Second, despite the scale of this study, we evaluated only a small fraction of the available neuroimaging and cognitive neuroscience literature. Broadening these evaluations will require contributions from the wider vision community. To facilitate this effort, we will release code and an accompanying website that allow researchers to benchmark their own datasets, and incorporate new experiments into the framework. Finally, our framework inherits an important limitation of the literature it evaluates. Although many classical findings were reproduced by current models, the interpretation of model failures remains uncertain. A failure may reflect a missing computation in the model, limitations of the original experiment, or an effect that is itself difficult to reproduce. Distinguishing among these possibilities will require closer integration between model evaluation and experimental replication. In the long term, model-based benchmarks may also help identify which findings are the most robust and which warrant further scrutiny.

In summary, our results show that current ANN models capture important aspects of the computations underlying category-selective cortex, but also reveal clear computational gaps. These strengths and limitations became apparent only by evaluating models across independent datasets and decades of cognitive neuroscience experiments within a common zero-shot framework. Together, these results provide a clearer picture of what current models have and have not learned about visual representations in category-selective brain regions. We hope this work encourages a shift toward evaluating models by how well they generalize across datasets and experiments, rather than by how accurately they predict randomly held-out images within a single dataset.

## Methods

Our goal was to determine how well ANN-based predictive (encoding) models generalize beyond the data used to build them (mapping data), and to identify the limits of their generalization abilities. We used a scaled-up across-dataset zero-shot evaluation framework with two complementary tests. *Prediction Tests* (Part 1) measured how accurately fixed encoding models predicted FFA, PPA, and EBA responses across six independent fMRI datasets. *Cognitive (neuroscience) tests* (Part 2) measured whether the same models reproduced 31 findings from 10 published cognitive neuroscience studies. Encoding models were developed using the standard approach ([16, 86] by learning a mapping from pretrained ANN features to fMRI responses using one of two mapping datasets (the Natural Scenes Dataset (NSD) and Murty185), and were then evaluated as is without further fitting (zero-shot). Additional analyses examined how generalization depended on stimulus distribution shift, brain region, mapping dataset, and ANN model family. For readability, the Methods follow the order of the Results, with each section providing the details needed to reproduce the corresponding analysis.

### ANN-based encoding models

#### Mapping fMRI datasets

We used two independent fMRI datasets to test the robustness of the model-to-brain mappings and its effects on zero-shot generalization (**Fig.3**).

#### Natural Scenes Dataset (NSD) [1]

To obtain a large and diverse set of mapping images, we used the 1000 natural scenes shared across NSD participants 1, 2, 5, and 7 (NSD-1000 subset) to fit the encoding models. Each image was presented three times to each participant. We used version 3 single-trial beta estimates (betas_fithrf_GLMdenoise_RR). Voxel responses were z-scored within each scanning session and then averaged across the three presentations of each stimulus. For the explicit comparison of within-versus across-dataset evaluation in **Fig.3a**, we fit models using the 800 images from the NSD-1000 subset and evaluated them on the remaining 200 images. FFA, PPA, and EBA were defined separately in each participant using an functional localizer reported in NSD at contrast threshold *t >* 7.

#### Murty185 [63]

Murty185 is a much smaller mapping dataset, containing fMRI responses from 4 participants to 185 naturalistic images sampled mostly from the THINGS [34] dataset. Each image was repeated median 20 times, has high-reliability response estimates and have previously been shown to support accurate encoding models of FFA, PPA, and EBA [63]. As in NSD, FFA, PPA, and EBA were independently localized in each participant (contrast threashold *t >* 7).

#### Model architectures

We trained 127 encoding models spanning different architectures including convolutional networks (CNNs), Vision Transformers, and recurrent CORnet architectures. Of these, 110 used pre-trained visual backbones that vary in training objective (supervised, self-supervised, vision–language, topographic, adversarially or blur-robust, and multi-task Taskonomy encoders) and in scale (parameter count and pre-training dataset size). As untrained controls, we included 13 randomly initialized copies of selected architectures. We additionally evaluated three V1-based VONENETs, which prepend a fixed Gabor-like primary-visual-cortex front-end to AlexNet, CORnet-S, or ResNet-50, and a pixels control whose backbone is an identity map on resized, ImageNet-normalized pixel intensities (no learned visual features). Each encoding model consisted of two components: a frozen backbone and a trainable mapper that transforms intermediate backbone features into predicted fMRI voxel responses. Backbone weights remained fixed throughout training. Detailed family labels and best layers info for each ROI are listed in Supplementary **Tab.S4** and **Tab.S5**.

## Model training

### Layer selection

We identified the optimal layer for every model and fROI by training mappers on all intermediate layers of the backbone (convolutional layers for CNNs, linear layers for transformers). We used a 80 20 train-validation split, to find the model layer with the validation prediction score (measured as Pearson R between observed and predicted responses).

### Mapping model features to brain responses

Our objective was to map image representations from pre-trained ANN backbones to fMRI voxel responses. Because backbone representations are high-dimensional, we used a more regularized and parameter efficient readout inspired by Klindt et al. [43] that separates the mapping into spatial and feature components. We selected the mapper based on the backbone model architecture type—ConvMapper for CNNs and TransformerMapper for transformers.

For CNNs, representations have shape R*^B×C×H×W^*, where *B* is the batch size, *C* is the number of feature channels, and *H* and *W* are the spatial dimensions of the feature map. For Vision Transformers (ViTs), representations have shape R*^B×N×D^*, where *B* is the batch size, *N* is the number of patches (or tokens, including the CLS token), and *D* is the embedding dimension. Both representations must be mapped to fMRI responses of shape R*^B×N^*^voxels^, where *N*_voxels_ is the number of voxels in a given region of interest. While a regularized ridge regression could serve as the mapping, the differences in representation sizes across models yield highly variable parameter counts (many times prohibitively large). To facilitate large-scale hyperparameter sweeps, we developed two compact, architecture-specific mapping functions: ConvMapper and TransformerMapper.

ConvMapper. This mapper handles CNN representations through three operations: (1) a single convolutional layer reduces the channel dimension from *C* to *C^′^* and compresses spatial dimensions; (2) an einops.reduce operation ([66]) averages over the spatial dimensions *H* and *W*, yielding shape R*^B×C′^* ; (3) a linear layer projects to the final voxel predictions of shape R*^B×N^*^voxels^ .

TransformerMapper. This mapper reduces ViT representations to voxel predictions through a factorized projection. Given a targ*_√_*e t hidden size *H*, it first factorizes *H* into two integers *p* and *q* with *p × q* = *H*, chosen to be as close to *H* as possible so that the factorization is near-square. These two integers become the reduced embedding and sequence sizes. Starting from an input of shape R*^B×N×D^*, the mapper applies two linear projections along different axes in sequence: the first compresses the embedding dimension from *D* to *p*, and the second compresses the sequence dimension from *N* to *q*, yielding a tensor of shape R*^B×p×q^*. This is flattened to R*^B×H^* and passed through a final linear layer to produce voxel predictions in R*^B×N^*^voxels^ . Because the two projections act on separate axes, this factorization mixes local (embedding) and global (patch-level) information at low cost while keeping the architecture compact. This mapping process results in comparable or better prediction performance than ridge regression (see also [2] but is significantly parameter efficient and can be scaled with GPU acceleration.

### Hyperparameter optimization

After selecting the readout layer, we performed a grid-search over the linear mapper’s hyperparameters for that layer, independently for each backbone × ROI. We used Adam with no learning-rate schedule, trained for 50 epochs with mean-squared error (MSE) loss, and split NSD 80/20 (seed 0) into training and validation sets. The grid comprised 20 log-spaced learning rates from 10*^−^*^5^ to 10*^−^*^1^ (np.logspace(-5, -1, 20)), batch sizes 16, 32, 48, 64, and three log-spaced weight-decay values 10*^−^*^7^, 10*^−^*^5^, 10*^−^*^3^, for a total of 240 configurations per backbone × ROI. Weights were checkpointed at the epoch with lowest validation MSE; among completed runs we retained the configuration with the highest validation Pearson correlation between flattened predicted and true voxel responses. That setting was used to retrain and save the final encoding model. The pixels control used a reduced grid (learning rate 10*^−^*^5^, batch size 32, weight decay 10*^−^*^7^).

#### Comparison between zero-shot evaluation and standard cross-validation

All evaluations in the paper are strictly zero-shot (across-dataset). Standard within-dataset cross-validation was reported only for the comparison with zero-shot prediction in Section 1.3. For this analysis, NSD models were trained on 800 of the 1,000 NSD mapping images and evaluated on the remaining 200 held-out images (using a deterministic random seed (0)). To construct the matched across-dataset evaluation, models were trained on all 185 images from Murty mapping dataset and evaluated zero-shot on the same 200 held-out NSD images. Thus, the two evaluations used identical test images and differed only in whether the model-to-brain mapping was estimated from images within the same dataset or from an independent dataset.

### Prediction Tests

To evaluate how well model predictions reproduce empirically observed brain responses zero-shot, we conducted both univariate and multivariate prediction tests.

#### Evaluation Datasets

We used six datasets exclusively for zero-shot evaluation. None of these datasets was used for model fitting, layer selection, or hyperparameter optimization. Together with the two mapping datasets described above (NSD and Murty185), this yielded eight fMRI datasets in total. When models were fit using one mapping dataset, the other mapping dataset could also serve as an independent zero-shot evaluation dataset, providing up to 7 evaluation datasets for each mapping configuration.

**BOLD5000v2** [13] contains BOLD responses from four participants while they viewed 4,196 natural images drawn from large-scale computer-vision image sets, providing a broad stimulus sample for benchmarking image-computable encoding models.

**King and Baker** [42] comprises fMRI responses from four participants to 4,916 natural images and was used to evaluate model predictions across a large, densely sampled image set optimized for representational analyses.

**Bonner and Epstein** [5] includes two independent stimulus sets, each containing 81 object images, with fMRI responses measured from five participants per set. These data provide a controlled test bed for evaluating object representations in high-level visual cortex.

**Wardle** [82] includes 96 stimuli, including 32 illusory faces in inanimate objects, 32 visually and semantically matched nonface objects, and 32 human faces. The fMRI experiment included 21 participants, of whom 16 were retained for analyses.

**BOLD Moments** [46] contains fMRI responses from ten participants to 102 dynamic movie clips sampled from the Moments in Time video dataset [54]. We used this dataset to evaluate model predictions for temporally extended, naturalistic visual input.

**NSD-Synthetic** [31] is a 7T fMRI dataset collected from the eight Natural Scenes Dataset participants while they viewed 284 controlled synthetic images. We used this dataset to test out-of-distribution generalization beyond natural-image training and evaluation regimes.

#### Dataset Ceiling

We estimated an inter-subject ceiling to characterize the consistency of neural responses within each dataset. We computed the corresponding response correlation for every pair of subjects and averaged across all subjects pairs. Thus, for a dataset with *S* subjects, the ceiling was defined as the mean pairwise inter-subject correlation, and was computed as:

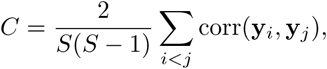

where y*_i_* and y*_j_* denote the response representations of participants *i* and *j*, respectively. Full inter-subject ceiling results across all datasets and ROIs are provided in **Table S2**.

#### Model Prediction Score

For the *univariate* prediction test, we asked whether models captured the voxel-averaged magnitude to images for each ROI. For each participant and ROI, we averaged the observed and predicted responses across voxels, yielding one observed and one predicted response for each stimulus. We then computed the Pearson correlation across stimuli between the predicted and observed response vectors and averaged the resulting correlations across participants. For the *multivariate* prediction test, we asked whether models captured the representational geometry within each ROI, following [63]. For each participant, we retained the full voxel-wise response pattern for every stimulus and constructed separate representational dissimilarity matrices (RDMs) from the observed and predicted responses [19, 45, 55]. Each RDM contained the pairwise Euclidean distance between the voxel-response patterns elicited by every pair of stimuli. We used Euclidean rather than correlation distance because differences in overall response magnitude carry meaningful information in category-selective regions and should be preserved [19, 81]. Model–brain correspondence was then measured as the Spearman correlation between the upper triangles of the predicted and observed RDMs and averaged across participants.

For the BOLD Moments dataset, stimuli were videos rather than static images. We uniformly sampled 10 frames from each video and independently passed each frame through the model. We then took the element-wise median of the predicted responses across frames, yielding a single predicted response for each video. The median reduced sensitivity to transient or atypical individual frames. These video-level predictions were evaluated against the observed fMRI responses using the same univariate and multivariate procedures described above.

The full heatmap of all 127 model performance on prediction tests is shown in **Fig.S1**. While **Fig.2c** showed a snapshot of Murty-mapped models performance, this full heatmap showed both mapping source with detailed model and evaluation dataset labels.

#### Quantifying zero-shot prediction difficulty

To quantify how difficult each evaluation dataset was for models to predict after mapping on NSD-1000, we measured how far model performance remained below the corresponding human inter-subject ceiling. For fairness, these dataset-level analyses included only the trained *110* models (and excludes pixel, randomly initialized, and V1-like baselines).

#### Normalized gap to ceiling

For each model *i*, evaluation dataset *d* and fROI *r*, we computed the model’s mean prediction score 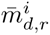 and normalized it by the corresponding mean inter-subject ceiling *c_d,r_*,

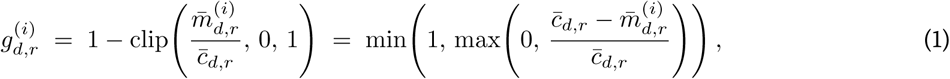

defined only when *c_d,r_* _>0_.

The normalized prediction gap ranges from 0 to 1, with 0 indicating that model performance reaches the inter-subject ceiling and larger values indicating worse prediction relative to that ceiling. Normalizing the predictions by the ceiling allowed model difficulty to be compared across datasets and ROIs with different levels of inter-subject reliability.

#### Wasserstein distance to the ceiling

As a complementary measure that considers the full distribution of prediction scores rather than only their means, we computed the first Wasserstein distance between the model-brain and human-human correlation distributions:

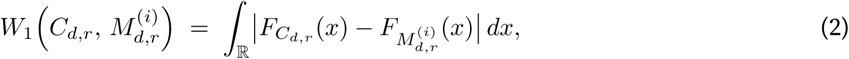

where *F* denotes the empirical cumulative distribution function (equal weight on each pairwise correlation). Larger Wasserstein distances indicate a greater separation between the model-brain and human-human correlation distributions.

#### Aggregation to dataset and fROI difficulty

To estimate the difficulty of each evaluation dataset, we first averaged the normalized prediction gap across the 110 trained models for each fROI, and then averaged across the fROIs available for that dataset. When comparing overall dataset difficulty, we additionally averaged across the univariate and multivariate analyses. To estimate the difficulty of each fROI, we averaged each model’s prediction gap across evaluation datasets and then summarized these values across models. Higher values indicate datasets or brain regions for which model predictions remained farther from the human ceiling.

#### Stimulus distance from the mapping dataset

To test whether models generalized less well to stimuli that differed from those used to build the encoding models, we measured the distance between each evaluation stimulus set and the NSD-1000 mapping images. Because there is no single definition of image similarity, we measured this distance in four different representation spaces. Pixel PCA captures low-level image similarity, DreamSim captures mid-level perceptual similarity [27], and AligNet [56] the high-level visual similarity between images. Images were embedded separately using each representation.

#### Pixel PCA

As a low-level baseline, images were resized to 224 × 224, flattened, and projected into a shared 1024-dimensional pixel space using randomized truncated SVD (TruncatedSVD, random_state=42) fit across all stimuli. This representation captures low-level pixel structure without relying on a learned visual model.

#### DreamSim

We embedded each image using the default pretrained DreamSim model and its standard preprocessing pipeline [27], yielding one perceptual feature vector per image.

#### AligNet

We embedded each image using the pretrained AligNet model [56], built on a DINOv2-B backbone, and used its triplet_logits output as the image representation.

For each representation, we measured how different the distribution of evaluation images was from the distribution of NSD images using squared Maximum Mean Discrepancy (MMD^2^, see also [49]). MMD compares all images in one dataset with all images in the other dataset and returns a single measure of how separated the two stimulus distributions are. Thus, a small MMD^2^ indicates that an evaluation dataset contains images similar to the NSD mapping images, whereas a large MMD^2^ indicates a larger stimulus distribution shift. For reproducibility, embeddings were *L*_2_-normalized before computing MMD^2^, and datasets containing more than 1,000 stimuli were randomly subsampled to 1000 images using a fixed random seed. We used an RBF kernel with bandwidth determined by the median cross-dataset distance. MMD^2^ was computed as:

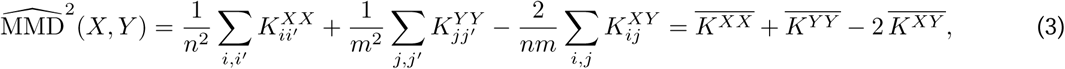

where *X* and *Y* are the NSD and evaluation embeddings, respectively, and *k* is the RBF kernel. We applied the same procedure independently in all four representation spaces and used the resulting NSD-to-dataset distances to test whether greater stimulus distribution shift predicted poorer zero-shot model generalization.

### Generalization Score

We quantified the relationship between stimulus distance and zero-shot generalization at two levels. Dataset-level generalization asks whether evaluation datasets farther from the NSD mapping images are harder for models overall. For each dataset, we averaged prediction difficulty across available fROIs and the 110 trained models, yielding one difficulty score per dataset. We then related these scores to the dataset’s MMD^2^ distance from NSD using ordinary least-squares (OLS) regression. A positive slope indicates that datasets farther from the mapping distribution are systematically harder for models to predict.

Model-level generalization instead asks how robustly each individual model transfers as evaluation stimuli move farther from the mapping distribution. For each model, we related its prediction difficulty across the seven evaluation datasets to their MMD^2^ distance from NSD and defined the resulting regression slope as that model’s generalization slope. Smaller slopes indicate models whose prediction performance degrades less with increasing stimulus distance and therefore generalize more robustly across datasets. Both analyses were performed separately for univariate and multivariate prediction tests and for the normalized prediction gap and Wasserstein-distance measures.

### Cognitive Tests

To test whether encoding models reproduce established functional properties of category-selective visual cortex, we assembled 31 experiments from 10 published fMRI studies. Studies were included when they (1) reported responses from individually localized FFA, PPA, or EBA, (2) made the original experimental stimuli publicly available, and (3) tested a specific hypothesis using controlled stimulus comparisons or manipulations. For each experiment, we identified the central reported effect and obtained the corresponding empirical responses from the original data when available. When numerical data were not publicly available, we digitized the reported responses from published figures using WebPlotDigitizer [67]. We then presented the original experimental stimuli to each fixed encoding model and asked whether its predicted responses reproduced the reported effect, without any additional fitting.

#### Published studies

The 31 experiments were drawn from ten studies spanning several established properties of category-selective cortex, including scene properties [3, 10, 36, 38, 58, 62], face and body configuration [9, 39], face pareidolia [82], and the relationship between neural representational geometry and visual-search behavior [15]. Individual experiments, target regions, stimulus conditions, and reported effects are detailed below.

**1. Rajimehr et al., 2011 [62] | Univariate | PPA.** Tested PPA responses to stimulus properties including spatial frequency and rectilinear versus curvilinear shapes.
**2. Kamps et al., 2019 [39] | Univariate | FFA, EBA.** Tested responses to faces with intact or scrambled internal and external facial features.
**3. Wardle et al., 2020 [82] | Multivariate | FFA, PPA.** Tested whether response patterns distinguish illusory faces from visually and semantically matched nonface objects.
**4. Park et al., 2015 [58] | Univariate | FFA, PPA.** Tested responses to indoor scenes parametrically varying in perceived spatial size and clutter.
**5. Bryan et al., 2016 [10] | Univariate | PPA.** Tested whether PPA scene selectivity persists when rectilinearity is controlled across scene and nonscene stimuli.
**6. Kamps et al., 2016 [38] | Univariate | PPA.** Tested responses to local scene elements and their arrangement within intact scenes.
**7. Cohen et al., 2017 [15] | Multivariate | FFA, PPA, EBA.** Tested the relationship between neural representational dissimilarity for object categories and human visual-search reaction times.
**8. Brandman and Yovel, 2016 [9] | Univariate | EBA.** Tested responses to whole bodies, isolated body parts, and configurations in which body parts were spatially rearranged.
**9. Berman et al., 2017 [3] | Univariate | PPA.** Tested scene-selective responses to images containing different ranges of spatial frequencies.
**10. Josephs and Konkle, 2020 [36] | Univariate | PPA.** Tested responses to objects, reachable-scale environments, and larger navigable scenes, including the effects of presenting multiple objects within reachable-scale environments.

### Quantifying replication of published findings

#### Human voting score

For experiments in which replication depended on the pattern of predicted responses, we obtained human judgments of replication from ten researchers with training background in computational neuroscience, none of whom were authors of published studies. This study was reviewed by the Institutional Review Board of the Georgia Institute of Technology.

Before viewing any model output, all ten raters attended a joint calibration session in which, for each experiment, they reviewed the hypothesis, experimental design, and original result and reached a shared understanding of the response pattern that would constitute a reproduction of the central finding. Raters then independently viewed the model-predicted results and judged, for each model, whether that pattern was reproduced (*yes*/*no*). Model identities were masked, and the order in which models were presented was randomized independently for each experiment and each rater. The ten raters were divided into two fixed groups of five. Within each experiment, one group evaluated the Murty-mapped models and the other evaluated the NSD-mapped models, and this assignment alternated across experiments. Consequently, although each model–experiment pair received five independent judgments, the ratings for each mapping dataset were, across the full set of experiments, contributed by all ten raters rather than by a single fixed group. We assessed agreement among raters using Spearman–Brown corrected split-half reliability. Across experiments, voting scores showed high reliability (mean *±* SD = 0.86 *±* 0.02).

For each model *×* experiment, we defined the voting score as:

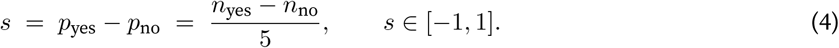

where *n*_yes_ and *n*_no_ are the numbers of raters judging the finding as reproduced or not reproduced, respectively. Scores range from 1, when all five raters judged the model to fail, to +1, when all five judged it to reproduce the finding. We averaged voting scores across experiments to obtain an overall replication score for each model, and across models to estimate the difficulty of each experiment. Higher scores indicate more consistent replication of the published finding.

### Statistically defined voting scores

14 experiments were scored directly using the statistical criterion from the original study, without human voting, because the replication outcome was unambiguous. Each experiment received +1 when the model predictions reproduced the reported statistical pattern and -1 otherwise.

### Validation against reported neural responses

As yet another independent validation, we extracted numerical responses from the original studies, using published data when available and WebPlotDigitizer when necessary [67]. We computed the mean squared error (MSE) between the reported and model-predicted response patterns and tested whether models receiving higher voting scores also produced lower MSE.

A full list of experiments and available quantify methods for each experiment is shown in **Table S3**. The full heatmap of all 127 model performance on cognitive tests is shown in **Fig.S2**.

## Supplementary

### Supplemental Figures

**Figure S1.**
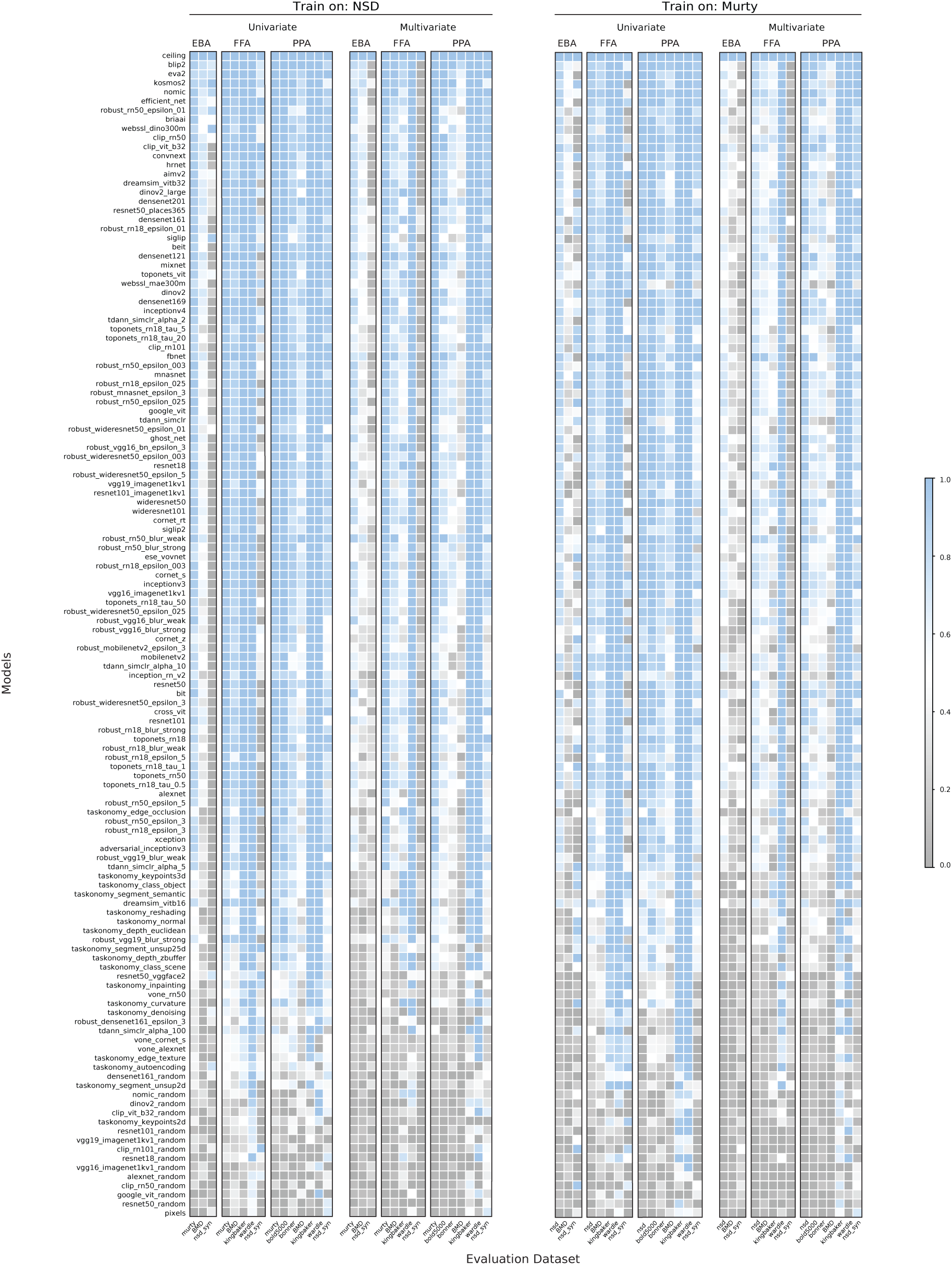
Zero-shot prediction across datasets for all 127 models. Each row corresponds to one ANN model and each column to one brain region and dataset. Models are ordered by decreasing prediction performance of NSD-mapped models. Scores are normalized by the human-to-human similarity ceiling, allowing comparison across datasets.

**Figure S2.**
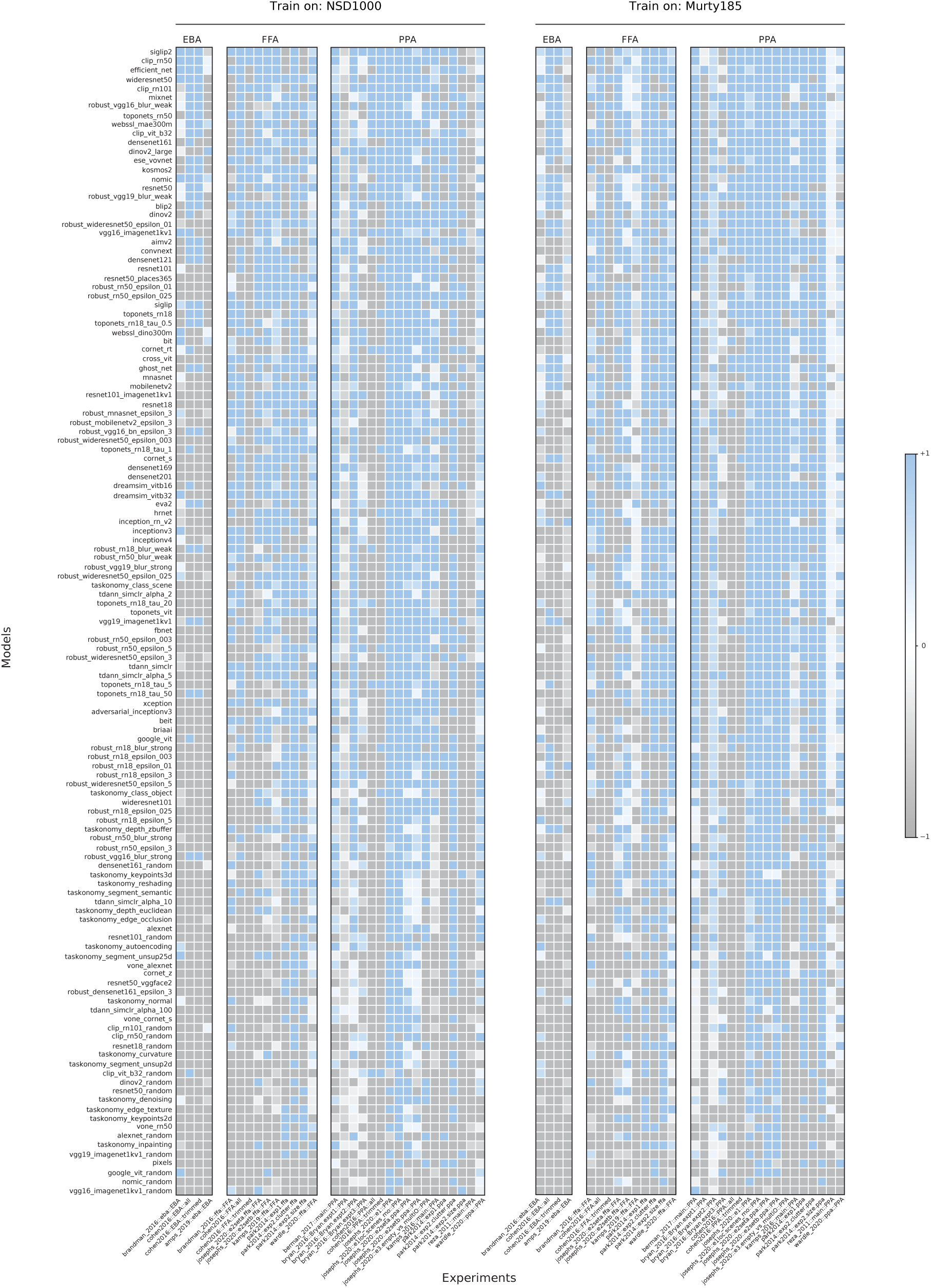
Zero-shot cognitive tests across experiments for all 127 models. Voting scores for PPA, FFA, and EBA across all models and 31 experiments from 10 papers. Models are ordered by decreasing prediction performance of NSD-mapped models. Each row corresponds to one ANN model and each column to one brain region and experiment.

**Figure S3.**
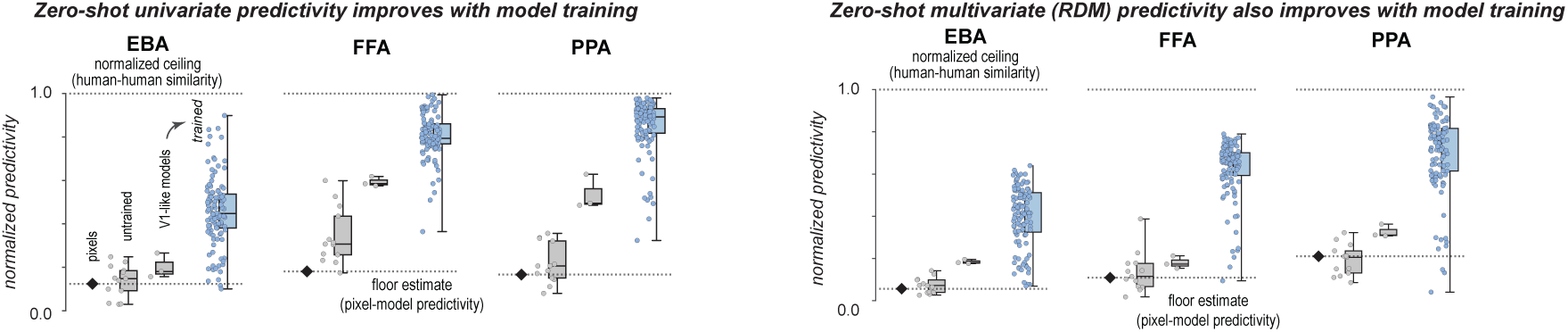
Zero-shot univariate and multivariate (RDM) prediction improves with model training (NSD-mapped models). Normalized univariate and multivariate prediction scores for EBA, FFA, and PPA comparing pixel-level predictions, untrained models, v1-like models, and trained ANN models. Each point represents one ANN model; boxes summarize the distribution across models. Dashed lines indicate the human-to-human similarity ceiling (top) and pixel-level prediction baseline (bottom). This figure shows results from NSD-mapped models on the 7 other independent datasets.

**Figure S4.**
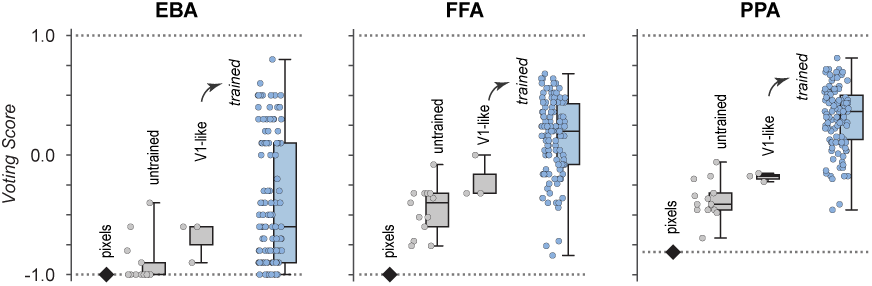
Zero-shot cognitive tests performance (voting score) improves with model training (Murty-mapped models). Voting scores for EBA, FFA, and PPA comparing pixel-level predictions, untrained models, v1-like models, and trained ANN models. Each point represents one ANN model; boxes summarize the distribution across models. This figure shows results from Murty-mapped models on 31 cognitive experiments.

**Figure S5.**
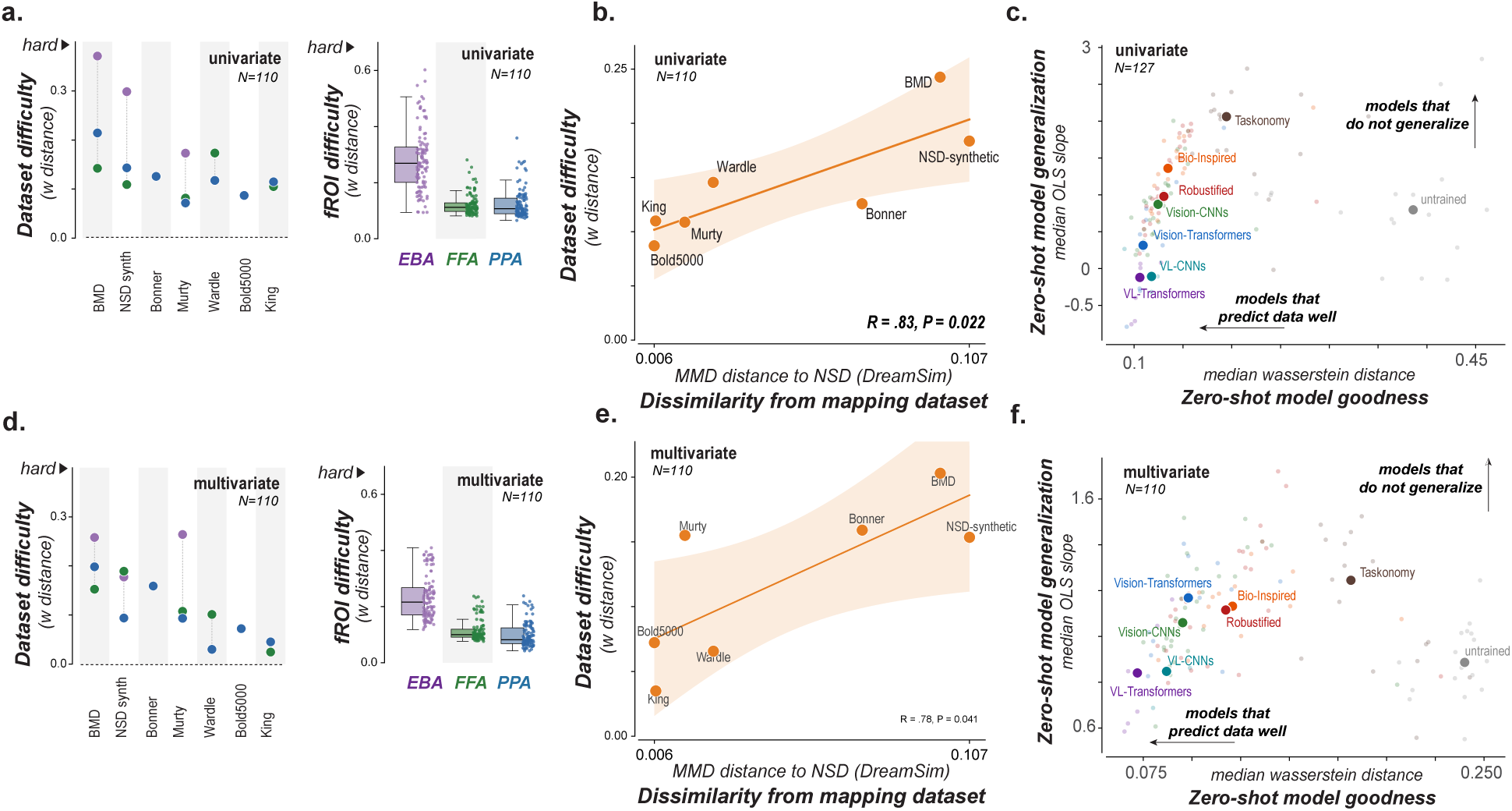
Zero-shot evaluation exposes structured limits to neural prediction, with prediction difficulty measured as wasserstein distance. **a.** Prediction difficulty varies systematically across datasets and brain regions. *Left*, univariate prediction difficulty for each evaluated fMRI dataset, quantified as the wasserstein distance between model-brain and human-human correlation distributions. Colors indicate the corresponding fROI. *Right*, fROI difficulty (y-axis) measured as wasserstein distance to ceiling across datasets for univariate data. **b.** Dataset difficulty increases with dissimilarity from the mapping dataset. Zero-shot univariate prediction difficulty plotted against the visual dissimilarity (DreamSim MMD) between each evaluation dataset and the mapping dataset (NSD). Prediction accuracy declined systematically as evaluation datasets became more dissimilar from the mapping images. **c.** Zero-shot evaluation identifies specific model families that both predict well and generalize well across datasets. Each point represents one ANN model. Zero-shot prediction performance (median wasserstein distance, x-axis), versus zero-shot generalization across datasets (slope of prediction performance across evaluation datasets, y-axis). Models in the bottom-left simultaneously achieve strong prediction accuracy and strong generalization. **d.** Same analysis as in **a.** using multivariate (RDM) prediction scores. **e.** Same analysis as in **b.** using multivariate (RDM) prediction scores. **f.** Same analysis as in **c.** using multivariate (RDM) prediction scores.

**Figure S6.**
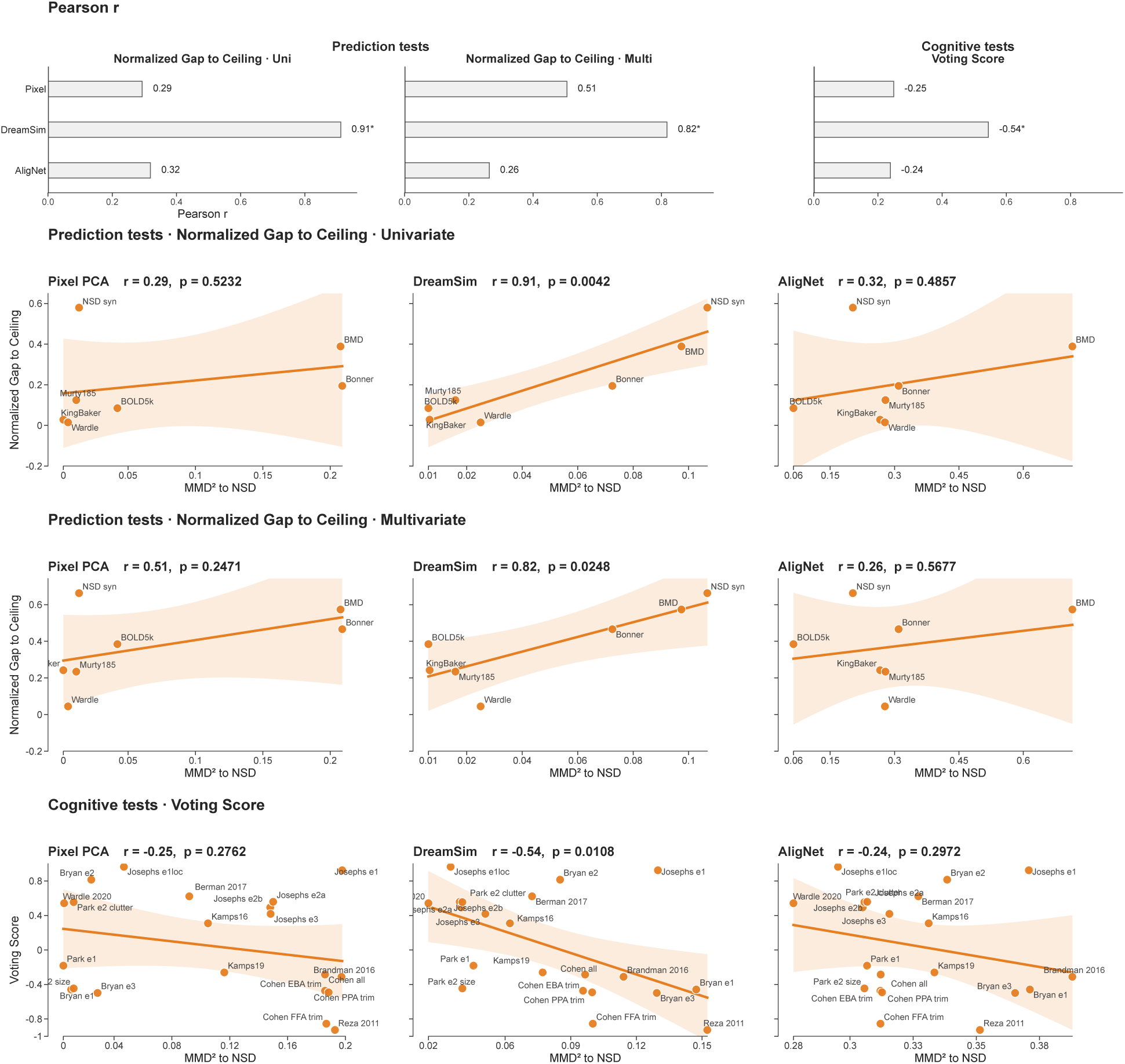
**Ordinary Least Squares (OLS) of zero-shot prediction tests and cognitive tests difficulty against stimuli dissimilarity to mapping dataset (NSD)**, for three embedding (*Pixel PCA*, *DreamSim*, *AligNet*). Each point is mean across 110-trained-model performance for one stimulus set (one fMRI dataset, or one cognitive experiment). The top row reports Pearson *r* for each outcome. *x* is RBF MMD^2^ to NSD (larger *x* = more dissimilar). Rows 1–4: *y* is prediction tests difficulty, quantified by normalized gap to ceiling or Wasserstein distance to ceiling correlation distributions (univariate and multivariate) for models trained from NSD. Bottom row: *y* is cognitive tests difficulty versus MMD^2^ on the 31 experiments, quantified by voting score. Before OLS, ROI-level results that used the same stimulus set are averaged. Shaded bands are 95% CI.

**Figure S7.**
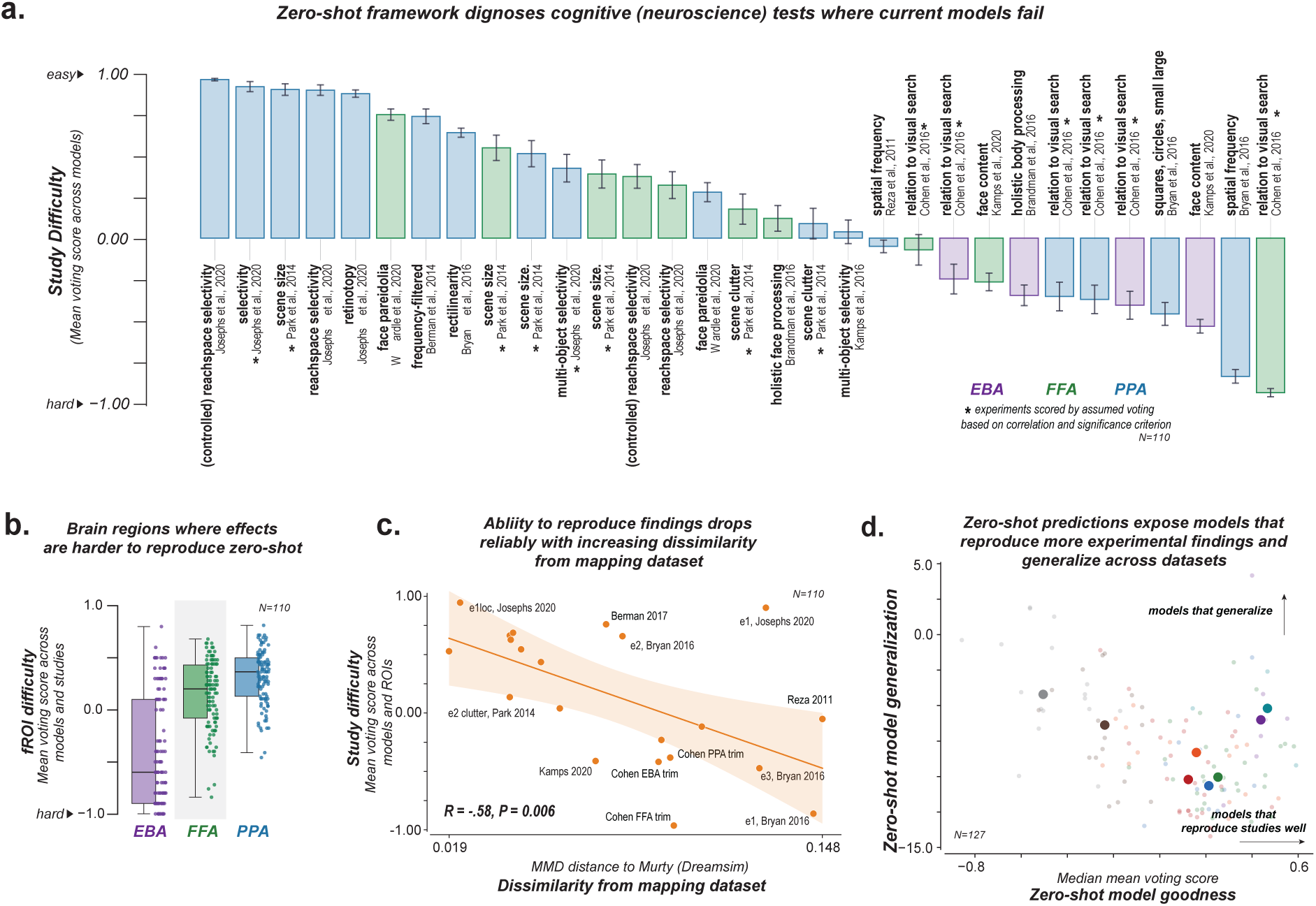
Understanding the challenges of model explanation on studies (Murty-mapped models). **a.** Zero-shot evaluation identifies cognitive neuroscience findings that current models reproduce and those they consistently fail to capture. Thirty-one cognitive neuroscience experiments are ranked by replication success (y-axis; voting score), from easiest (left) to hardest (right). Colors indicate the corresponding category-selective region (EBA, FFA, or PPA). Positive scores indicate successful replication of the published finding, whereas negative scores indicate systematic failures. **b.** Brain regions differ in the difficulty of reproducing cognitive neuroscience findings. Replication success (voting score, y-axis) summarized across experiments for EBA, FFA, and PPA. Each point represents one model; boxes summarize the distribution across experiments. **c.** Experiments become harder to reproduce as they become more dissimilar from the mapping dataset. Replication success is plotted against the visual dissimilarity (DreamSim MMD) between each cognitive neuroscience experiment and the mapping dataset **d.** Models that generalize across datasets also reproduce more cognitive neuroscience findings. Each point represents one ANN model. The x-axis shows overall replication success across the 31 cognitive neuroscience experiments, and the y-axis quantifies zero-shot generalization across experiments (median OLS slope). Models in the upper-right reproduce the largest fraction of cognitive neuroscience findings and generalizing consistently across experiments.

### Supplemental Tables

**Table S1.** Paired comparisons of NSD-versus Murty-mapped models zero-shot prediction performance on six held-out datasets. Each test is across *N* =110 trained models (df=109). Positive *t* indicates higher accuracy for NSD-mapped models. Overall averages pool all available dataset*×*ROI scores per model.

| Analysis | Dataset | $t(109)$ | $p$ | Mean $\Delta$<br>(NSD–Murty) | ROIs |
| --- | --- | --- | --- | --- | --- |
| Univariate | **Overall** | **4.91** | **$3.20 \times 10^{-6}$** | 0.015 | all |
| | BOLD5000 | 6.50 | $2.53 \times 10^{-9}$ | 0.030 | PPA |
| | Bonner 2021 | 2.78 | $6.46 \times 10^{-3}$ | 0.026 | PPA |
| | King & Baker 2019 | 2.88 | $4.75 \times 10^{-3}$ | 0.005 | FFA, PPA |
|  | Wardle 2020 | −0.89 | 0.373 | −0.002 | FFA, PPA |
|  | **BMD 2024** | **13.76** | **$1.37 \times 10^{-25}$** | 0.069 | EBA, FFA, PPA |
| | NSD-synthetic | −3.07 | $2.69 \times 10^{-3}$ | −0.028 | EBA, FFA, PPA |
| Multivariate | **Overall** | **10.66** | **$1.31 \times 10^{-18}$** | 0.012 | all |
| | BOLD5000 | 7.57 | $1.30 \times 10^{-11}$ | 0.016 | PPA |
|  | Bonner 2021 | −0.07 | 0.943 | −0.000 | PPA |
|  | King & Baker 2019 | 1.37 | 0.172 | 0.002 | FFA, PPA |
|  | Wardle 2020 | 1.43 | 0.157 | 0.001 | FFA, PPA |
|  | **BMD 2024** | **11.70** | **$5.66 \times 10^{-21}$** | 0.037 | EBA, FFA, PPA |
|  | NSD-synthetic | 1.84 | 0.069 | 0.005 | EBA, FFA, PPA |

**Table S2.** Inter-subject ceiling means used to characterize the consistency of neural responses within each dataset. Values report the mean ceiling correlation across subject pairs for each dataset and ROI, shown separately for univariate and multivariate analyses. *N* is the number of subjects.

| Dataset | $N$ | ROI | Ceiling Mean (Univariate) | Ceiling Mean (Multivariate) | Dataset Type |
| --- | --- | --- | --- | --- | --- |
| bmd_2024 | 10 | EBA | 0.6595 | 0.4028 | Video |
|  |  | FFA | 0.6667 | 0.3987 |  |
|  |  | PPA | 0.5369 | 0.2804 |  |
| bold_5000 | 3 | PPA | 0.4169 | 0.1829 | Natural images |
| bonner_2021 | 4 | PPA | 0.6303 | 0.3397 | Natural images |
| kingbaker_2019 | 5 | FFA | 0.2898 | 0.0489 | Natural images |
|  |  | PPA | 0.1423 | 0.0685 |  |
| murty185 | 4 | EBA | 0.8452 | 0.6627 | Natural images |
|  |  | FFA | 0.8201 | 0.5756 |  |
|  |  | PPA | 0.8141 | 0.5658 |  |
| nsd_1000 | 4 | EBA | 0.7443 | 0.5169 | Natural images |
|  |  | FFA | 0.7301 | 0.4637 |  |
|  |  | PPA | 0.7043 | 0.4007 |  |
| nsd_syn | 8 | EBA | 0.2536 | 0.2257 | Synthetic |
|  |  | FFA | 0.1409 | 0.1943 |  |
|  |  | PPA | 0.6041 | 0.2928 |  |
| wardle_2020 | 16 | FFA | 0.1706 | 0.0320 | Natural images |
|  |  | PPA | 0.0783 | 0.0082 |  |

**Table S3.** Experiment membership and mean scores by mapping dataset. Within each mapping block, experiments are **ranked by replication from easy to hard** (mean across *N* = 110 models; higher voting score = easier). MSE is also provided (lower = closer fit). Cells show the mean score when included, and –when excluded.

| Mapping dataset | Experiment (easy → hard) | ROI | Category properties | Human voting score (17 exp) | Statistically defined voting score (14 exp) | MSE (22 exp) |
| --- | --- | --- | --- | --- | --- | --- |
| NSD | josephs_2020, e1loc.scenes_mo | PPA | selectivity | – | 0.96 | 0.53 |
| NSD | josephs_2020, e1 | PPA | retinotopy | 0.92 | – | – |
| NSD | josephs_2020, e2seta.ppa | PPA | reachspace selectivity | 0.87 | – | 0.25 |
| NSD | bryan_2016, Bryan.expt2 | PPA | rectilinearity | 0.81 | – | 0.17 |
| NSD | josephs_2020, e2setb.ppa | PPA | (controlled) reachspace selectivity | 0.71 | – | 0.25 |
| NSD | berman_2017, main | PPA | frequency-filtered | 0.62 | – | 0.46 |
| NSD | park2014, exp2.clutter | FFA | scene clutter | – | 0.60 | 0.77 |
| NSD | wardle_2020, ffa | FFA | face pareidolia | 0.56 | – | – |
| NSD | wardle_2020, ppa | PPA | face pareidolia | 0.52 | – | – |
| NSD | park2014, exp2.clutter | PPA | scene clutter | – | 0.51 | 0.77 |
| NSD | josephs_2020, e3.empty_vs_multiO | PPA | multi-object selectivity | – | 0.42 | 0.39 |
| NSD | kamps_2020, ffa | FFA | face content | 0.35 | – | 0.55 |
| NSD | kamps_2016, main | PPA | multi-object selectivity | 0.31 | – | 0.34 |
| NSD | josephs_2020, e2setb.ffa | FFA | (controlled) reachspace selectivity | 0.28 | – | 0.64 |
| NSD | josephs_2020, e2seta.ffa | FFA | reachspace selectivity | 0.24 | – | 0.94 |
| NSD | cohen2016, all | FFA | relation to visual search | – | 0.13 | – |
| NSD | park2014, exp1 | FFA | scene size | – | 0.13 | 1.34 |
| NSD | brandman_2016, ffa | FFA | holistic face processing | -0.08 | – | 0.86 |
| NSD | park2014, exp2.size | FFA | scene size | – | -0.24 | 0.87 |
| NSD | cohen2016, all | EBA | relation to visual search | – | -0.35 | – |
| NSD | bryan_2016, Bryan.expt1 | PPA | squares, circles, small large | -0.46 | – | 1.40 |
| NSD | cohen2016, trimmed | EBA | relation to visual search | – | -0.47 | – |
| NSD | cohen2016, trimmed | PPA | relation to visual search | – | -0.49 | – |
| NSD | park2014, exp1 | PPA | scene size | – | -0.49 | 1.78 |
| NSD | bryan_2016, Bryan.expt3 | PPA | spatial frequency | -0.50 | – | 1.34 |
| NSD | brandman_2016, eba | EBA | holistic body processing | -0.55 | – | 1.90 |
| NSD | cohen2016, all | PPA | relation to visual search | – | -0.64 | – |
| NSD | park2014, exp2.size | PPA | scene size | – | -0.65 | 0.87 |
| NSD | cohen2016, trimmed | FFA | relation to visual search | – | -0.85 | – |
| NSD | kamps_2020, eba | EBA | face content | -0.87 | – | 1.01 |
| NSD | reza_2011, main | PPA | spatial frequency | -0.93 | – | 0.85 |
| Murty | josephs_2020, e2setb.ppa | PPA | (controlled) reachspace selectivity | 0.99 | – | 0.11 |
| Murty | josephs_2020, e1loc.scenes_mo | PPA | selectivity | – | 0.95 | 0.53 |
| Murty | park2014, exp2.size | PPA | scene size | – | 0.93 | 0.47 |
| Murty | josephs_2020, e2seta.ppa | PPA | reachspace selectivity | 0.92 | – | 0.17 |
| Murty | josephs_2020, e1 | PPA | retinotopy | 0.90 | – | – |
| Murty | wardle_2020, ffa | FFA | face pareidolia | 0.77 | – | – |
| Murty | berman_2017, main | PPA | frequency-filtered | 0.76 | – | 0.42 |
| Murty | bryan_2016, Bryan.expt2 | PPA | rectilinearity | 0.66 | – | 0.17 |
| Murty | park2014, exp1 | FFA | scene size | – | 0.56 | 0.84 |
| Murty | park2014, exp1 | PPA | scene size | – | 0.53 | 0.94 |
| Murty | josephs_2020, e3.empty_vs_multiO | PPA | multi-object selectivity | – | 0.44 | 0.43 |
| Murty | park2014, exp2.size | FFA | scene size | – | 0.40 | 0.47 |
| Murty | josephs_2020, e2setb.ffa | FFA | (controlled) reachspace selectivity | 0.39 | – | 0.34 |
| Murty | josephs_2020, e2seta.ffa | FFA | reachspace selectivity | 0.33 | – | 0.49 |
| Murty | wardle_2020, ppa | PPA | face pareidolia | 0.29 | – | – |
| Murty | park2014, exp2.clutter | FFA | scene clutter | – | 0.18 | 1.21 |
| Murty | brandman_2016, ffa | FFA | holistic face processing | 0.12 | – | 0.80 |
| Murty | park2014, exp2.clutter | PPA | scene clutter | – | 0.09 | 1.21 |
| Murty | kamps_2016, main | PPA | multi-object selectivity | 0.04 | – | 0.67 |
| Murty | reza_2011, main | PPA | spatial frequency | -0.05 | – | 0.74 |
| Murty | cohen2016, all | FFA | relation to visual search | – | -0.07 | – |
| Murty | cohen2016, all | EBA | relation to visual search | – | -0.25 | – |
| Murty | kamps_2020, ffa | FFA | face content | -0.27 | – | 0.60 |
| Murty | brandman_2016, eba | EBA | holistic body processing | -0.36 | – | 1.37 |
| Murty | cohen2016, all | PPA | relation to visual search | – | -0.36 | – |
| Murty | cohen2016, trimmed | PPA | relation to visual search | – | -0.38 | – |
| Murty | cohen2016, trimmed | EBA | relation to visual search | – | -0.42 | – |
| Murty | bryan_2016, Bryan.expt3 | PPA | spatial frequency | -0.47 | – | 1.00 |
| Murty | kamps_2020, eba | EBA | face content | -0.55 | – | 1.05 |
| Murty | bryan_2016, Bryan.expt1 | PPA | squares, circles, small large | -0.86 | – | 2.13 |
| Murty | cohen2016, trimmed | FFA | relation to visual search | – | -0.96 | – |

**Table S4.**
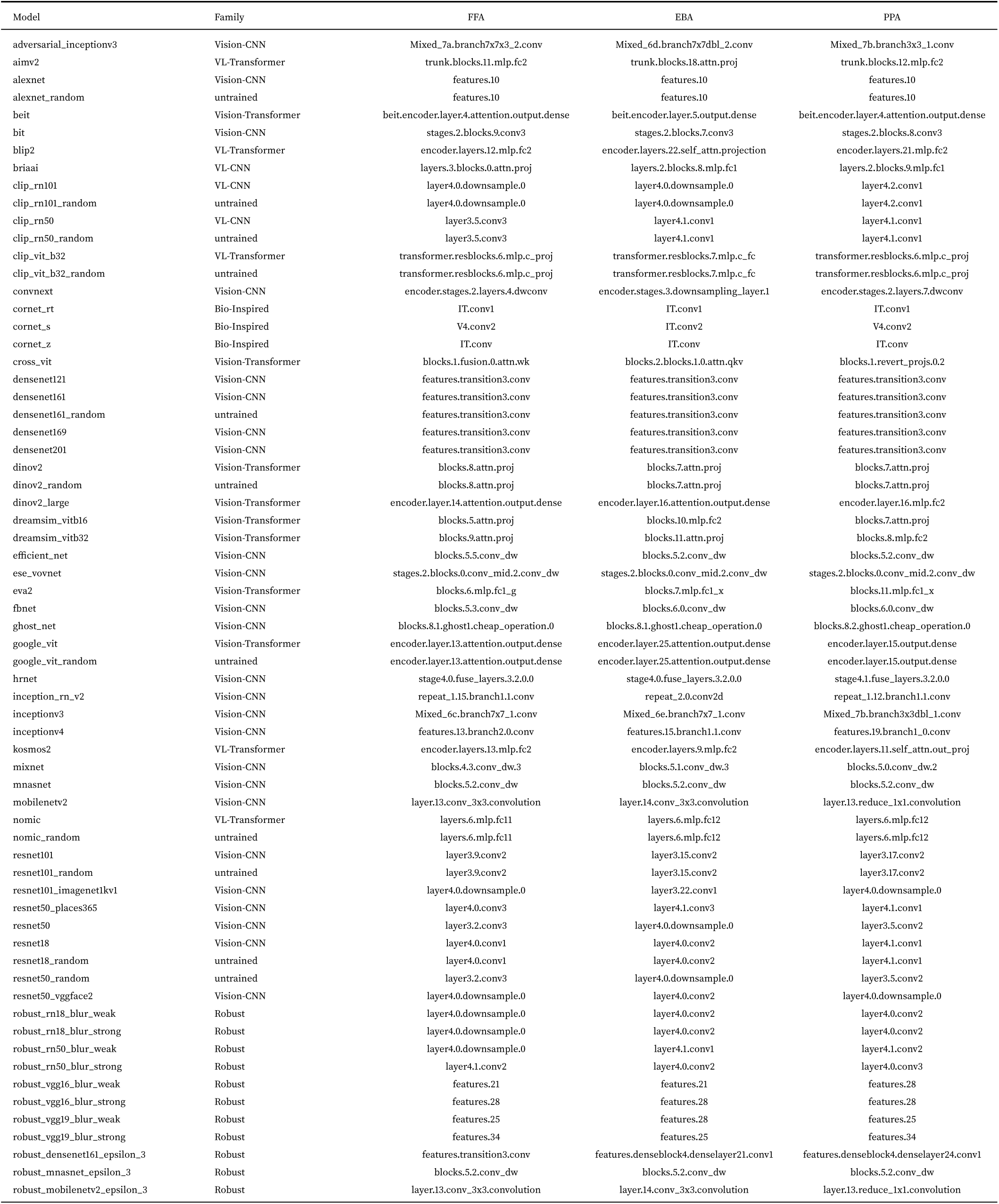
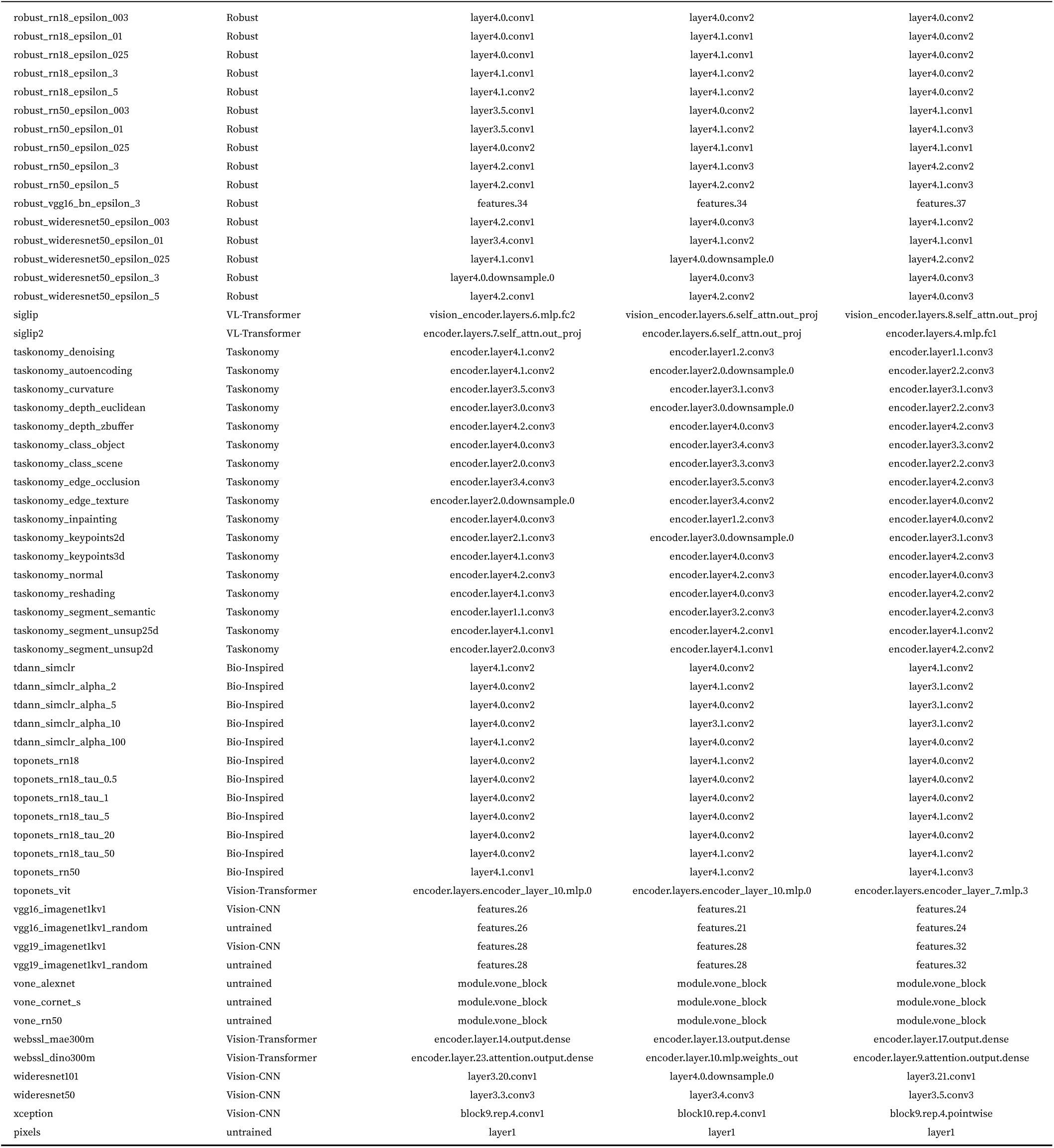
Models mapped from Murty. Family follows the model taxonomy used throughout (VL-Transformer, VL-CNN, Vision-Transformer, Vision-CNN, Bio-Inspired, Robust, Taskonomy, untrained).

**Table S5.**
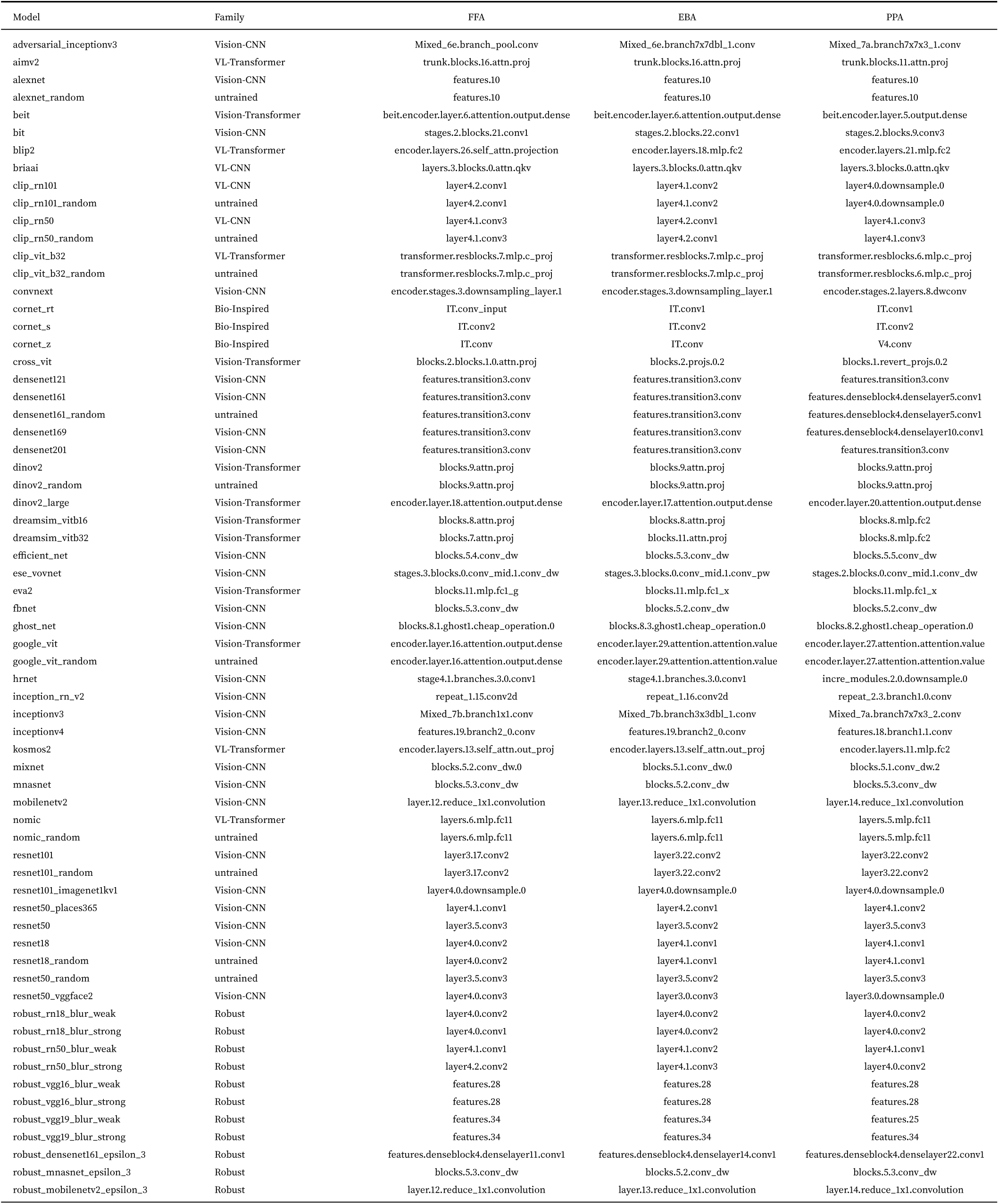
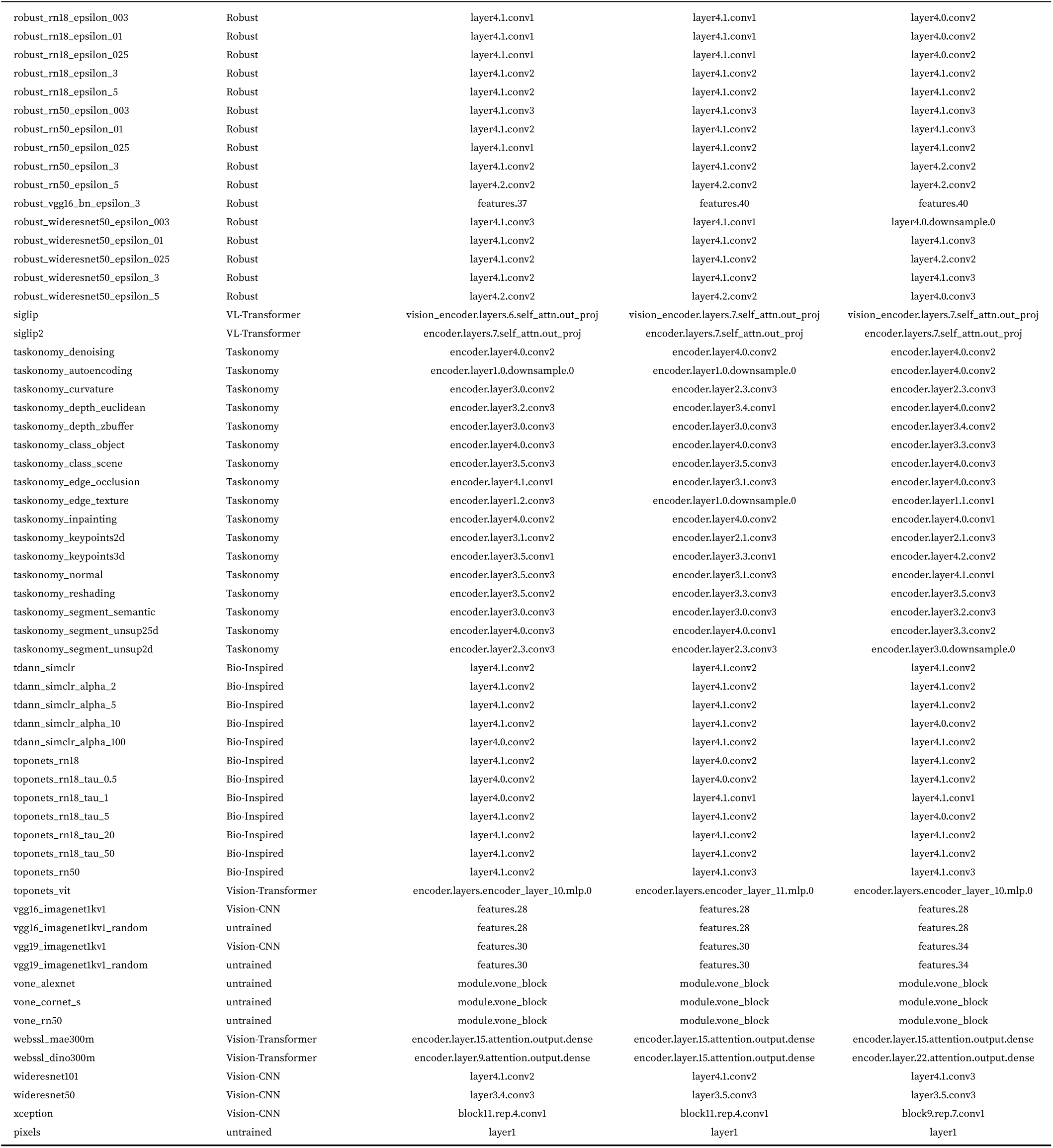
Models mapped from NSD. Family follows the model taxonomy used throughout (VL-Transformer, VL-CNN, Vision-Transformer, Vision-CNN, Bio-Inspired, Robust, Taskonomy, untrained).

## Acknowledgments

This work was supported by the NIH Pathway to Independence Award (R00EY032603), the NSF Nexus Computational Support program (Allocation No. SOC250049), and startup funds from Georgia Tech. We thank Michael Cohen for providing feedback on the paper and members of the Vision, Cognition, and Computation Lab for helpful discussions and feedback at multiple stages of this work.

## Author Contributions

N.A.R.M, N.K, A.A., R.W. and M.D conceived the study. R.W. led the model evaluation pipeline and M.D. led the model training pipeline. A.A. developed the initial prototype of the study replication pipeline, which was further developed by R.W. A.D. performed fMRI dataset preprocessing. R.W., M.D., S.C., K.R., K.D. and R.K. trained the models, and R.W., S.C., K.R. and K.D. configured the datasets and experiments. A.D., H.A. and R.K. performed the model categorization. A.D. and H.F. contributed to the evaluation protocol. E.M. extracted data from published studies. R.W. analysed the evaluation data and interpreted the results with N.A.R.M. R.W., Y.L., and N.A.R.M. prepared the figures. R.W., M.D., N.K. and N.A.R.M. wrote the manuscript. N.A.R.M. supervised the project. All authors discussed the results and edited the manuscript.

